# Flexible gating between subspaces in a neural network model of internally guided task switching

**DOI:** 10.1101/2023.08.15.553375

**Authors:** Yue Liu, Xiao-Jing Wang

**Affiliations:** New York University

## Abstract

Behavioral flexibility relies on the brain’s ability to switch rapidly between multiple tasks, even when the task rule is not explicitly cued but must be inferred through trial and error. The underlying neural circuit mechanism remains poorly understood. We investigated recurrent neural networks (RNNs) trained to perform an analog of the classic Wisconsin Card Sorting Test. The networks consist of two modules responsible for rule representation and sensorimotor mapping, respectively, where each module is comprised of a circuit with excitatory neurons and three major types of inhibitory neurons. We found that rule representation by self-sustained persistent activity across trials, error monitoring and gated sensorimotor mapping emerged from training. Systematic dissection of trained RNNs revealed a detailed circuit mechanism that is consistent across networks trained with different hyperparameters. The networks’ dynamical trajectories for different rules resided in separate subspaces of population activity; the subspaces collapsed and performance was reduced to chance level when dendrite-targeting somatostatin-expressing interneurons were silenced, illustrating how a phenomenological description of representational subspaces is explained by a specific circuit mechanism.

## Introduction

A signature of cognitive flexibility is the ability to adapt to a changing task demand. Oftentimes, the relevant task is not explicitly instructed, but needs to be inferred from previous experiences. In laboratory studies, this behavioral flexibility is termed un-cued task switching. A classic task to evaluate this ability is the Wisconsin Card Sorting Test (WCST) [1]. During this task, subjects are presented with an array of cards, each with multiple features, and should respond based on the relevant feature dimension (i.e. the task rule) that changes across trials. Crucially, subjects are not instructed on when the rule changes, but must infer the currently relevant rule based on the outcome of previous trials. This is different from cued rule switching paradigms where an external cue provides information to the subjects about the relevant rule for each trial (e.g. [2, 3]). Intact performance on un-cued task switching depends on higher-order cortical areas such as the prefrontal cortex (PFC) [4, 5, 6, 7, 8], which has been proposed to represent the task rule and modulate the activity of other cortical areas along the sensorimotor pathway [9].

Four essential neural computations must be implemented by the neural circuitry underlying un-cued task switching. First, it should maintain an internal representation of the task rule across multiple trials when the rule is unchanged. Second, soon after the rule switches, the animal will inevitably make errors and receive negative feedback, since the switches are un-cued. This negative feedback should induce an update to the internal representation of the task rule. Third, the neural signal about the task rule should be communicated to the brain regions responsible for sensory processing and action selection. Fourth, this rule signal should be integrated with the incoming sensory stimulus to produce the correct action. In contrast, in cued rule switching tasks, the first two computations (rule maintenance and rule updating) are not required, since an external cue about the rule is provided during every trial.

Prior work has identified neural correlates of cognitive variables presumed to underlie these computations including rule [10], feedback [10, 11, 12] and conjunctive codes for sensory, rule, and motor information [13]. In addition, different types of inhibitory neurons are known to play different functional roles in neural computation: while parvalbumin (PV)- expressing interneurons are suggested to underlie feedforward inhibition [14], interneurons that express somatostatin (SST) and vasoactive intestinal peptide (VIP) have been proposed to mediate top-down control [15, 16, 17, 18]. In particular, SST and VIP neurons form a disinhibitory motif [19, 20, 21] that has been hypothesized to instantiate a gating mechanism for flexible routing of information in the brain [22]. However, there is currently a lack of mechanistic understanding of how these neural representations and cell-type-specific mechanisms work together to accomplish un-cued task switching.

To this end, we used computational modeling to formalize and discover mechanistic hypotheses. In particular, we used tools from machine learning to train a collection of biologically-informed recurrent neural networks (RNNs) to perform an analog of the WCST used in monkeys [23, 10, 11]. Training RNN [24] does not presume a particular circuit solution, enabling us to explore potential mechanisms. For this purpose, it is crucial that the model is biologically plausible. To that end, each RNN was set up to have two modules: a “PFC” module for rule maintenance and switching and a “sensorimotor” module for executing the sensorimotor transformation conditioned on the rule. To explore the potential functions of different neuronal types in this task, each module of our network consists of excitatory neurons with somatic and dendritic compartments as well as PV, SST and VIP inhibitory neurons, where the connectivity between cell types is constrained by experimental data (Methods).

After training, we performed extensive dissection of the trained models to reveal that close interplay between local and across-area processing was essential for solving the WCST. First, we found that abstract cognitive variables were distinctly represented in the PFC module. In particular, two subpopulations of excitatory neurons emerge in the PFC module - one encodes the task rule and the other shows mixed-selectivity that nonlinearly depends on rule and negative feedback. Notably, neurons with similar response profiles have been reported in neurophysiological recordings of monkeys performing the same task [10, 11]. Second, we identified interesting structures in the local connectivity between different neuronal assemblies within the PFC module, which enabled us to compress the high-dimensional PFC module down to a low-dimensional simplified network. Importantly, the neural mechanism for maintaining and switching rule representation is readily interpretable in the simplified network. Third, we found that the rule information in the PFC module is communicated to the sensorimotor module via structured long-range connectivity patterns along the monosynaptic excitatory pathway, the di-synaptic pathway that involves PV neurons, as well as the trisynaptic disinhibitory pathway that involves SST and VIP neurons. In addition, different dendritic branches of the same excitatory neuron in the sensorimotor module can be differentially modulated by the task rule depending on the sparsity of the local connections from the dendrite-targeting SST neurons. Fourth, single neurons in the sensorimotor module show nonlinear mixed selectivity to stimulus, rule and response, which crucially depends on the activity of the SST neurons. On the population level, the neural population activity in the sensorimotor module during different task rules occupy nearly orthogonal subspaces, which is disrupted by silencing the SST neurons. Lastly, we found structured patterns of input and output connections for the sensorimotor module, which enables appropriate rule-dependent action selection. These results are consistent across dozens of trained RNNs with different types of dendritic nonlinearities and initializations, therefore pointing to a common neural circuit mechanism underlying the WCST.

## Results

### Training modular recurrent neural networks with different types of inhibitory neurons

We trained a collection of modular RNNs to perform the WCST. Each RNN consists of two modules: the “PFC” module receives an input about the outcome of the previous trial, and was trained to output the current rule; the “sensorimotor” module receives the sensory input and was trained to generate the correct choice (Figure 1b). The inputs and outputs were represented by binary vectors (Figure 1b, Methods). Each module was endowed with excitatory neurons with somatic and two dendritic compartments, as well as three major types of genetically-defined inhibitory neurons (PV, SST and VIP). Different types of neurons have different inward and outward connectivity patterns constrained by experimental data in a binary fashion (Methods, Figure 1b). The somata of all neurons were modeled as standard leaky units with a rectified linear activation function. The activation of the dendritic compartments, which can be viewed as a proxy for the dendritic voltage, is a nonlinear sigmoidal function of the excitatory and inhibitory inputs they receive (Methods). The specific form of the nonlinearity is inspired by experiments showing that inhibition acts subtractively or divisively on the dendritic nonlinearity function depending on its relative location to the excitation along the dendritic branch [25]. Therefore, we trained a collection of RNNs, each with either subtractive or divisive dendritic nonlinearity, to explore the effect of dendritic nonlinearity on the network function.

**Figure 1:**
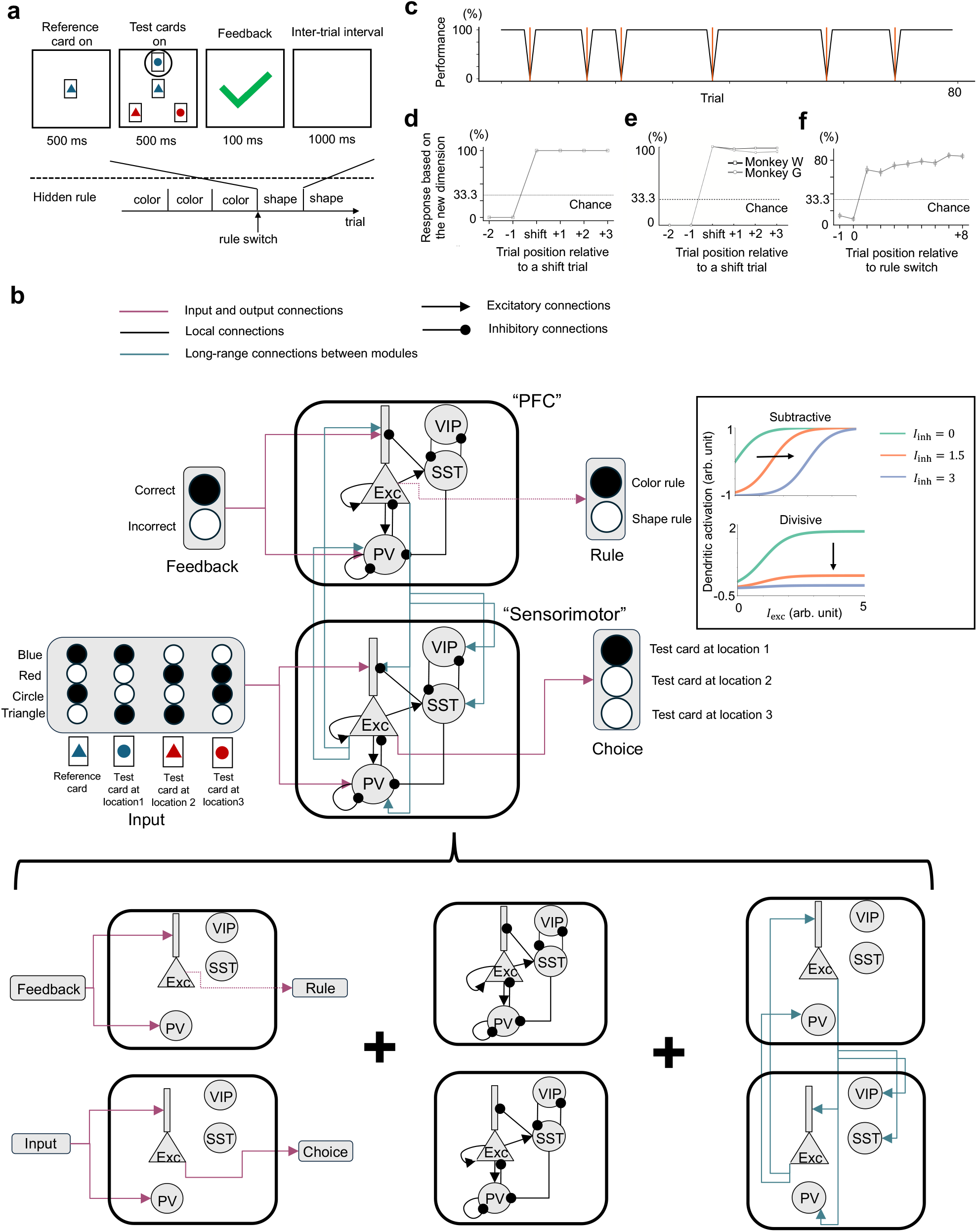
Model setup and performance. **a**. The schematic of the WCST task. Subjects are required to choose the card that matches the reference card at the center in either shape or color, depending on a hidden rule that switches after a number of trials. **b**. The RNN contains a “PFC” module and a “sensorimotor” module. The PFC module receives an input about the feedback of the previous trial, and was trained to produce the current rule. The sensorimotor module receives the sensory input and was trained to produce the correct choice. Each module is endowed with excitatory neurons and three types of inhibitory neurons: PV, SST and VIP. The cell-type-specific connectivity is constrained by experimental data (Methods). Bottom panel shows the decomposition of the model architecture into the input and output connectivity (left, magenta. The dashed line from PFC to rule represents the fact that the PFC module was trained to represent the rule but there are no explicit rule output neurons in the model), the local recurrent connectivity (middle, black) and inter-modular connectivity (green, right). Each excitatory neuron is modeled with a somatic and two dendritic compartments. For visual simplicity only one compartment was shown. Inset shows the relationship between the excitatory input onto the dendrite and the dendritic activity, for different levels of inhibitory inputs, as well as for the two types of dendritic nonlinearities used. Arb. units: arbitrary units; Exc: excitatory neurons; *I*_exc_: excitatory input; *I*_inh_: inhibitory input. **c**. The performance of the model during testing, for an example network. The network made one error after each rule switch (red vertical lines) and quickly recovered its performance. **d**. Performance as a function of trial position relative to the first correct trial after rule change, or the “shift” trial, for the same example network as in **c**. **e**. Performance of two monkeys while performing the same WCST analog as a function of trial position relative to the shift trial. Figure adapted from Ref. [28] (©2011 by the Massachusetts Institute of Technology. All rights reserved.) **f**. The performance of an example model where training was stopped before it reached perfect performance. This model exihibits more gradual switching between rules. Error bars represent standard error of the mean across *n* = 100 switches.

The task we trained the network on is a WCST variant used in monkey experiments [23, 10, 11, 8] (Figure 1a). During each trial, a reference card with a particular color and shape is presented on the screen for 500 ms, after which three test cards appear around the reference card for another 500 ms. Each card can have one of the two colors (red or blue) and one of the two shapes (square or triangle). A choice should be made for the location that contains the test card that has the same relevant feature (color or shape) as the reference card, after which the outcome of the trial is given, followed by an inter-trial interval. The relevant feature to focus on, or the task rule, changes randomly every few trials. Critically, the rule changes were not cued, requiring the network to memorize the rule of the last trial using its own dynamics. Therefore, the network dynamics should be carried over between consecutive trials, rather than reset at the end of each trial as has been done traditionally [2, 26]. To this end, the network operated continuously across multiple trials during training, and the loss function was aggregated across multiple trials (Methods). We used supervised learning to adjust the strength of all the connections (input, recurrent and output) by minimizing the mean squared error between the output of both modules and the desired output (rule for the PFC module and response for the sensorimotor module).

Notably, only the connections between certain cell types are non-zero and can be modified. This is achieved using a mask matrix, similar to [27] (Methods).

After training converged, we tested the models by running them continuously across 80 trials of WCST with rule switches at randomly chosen trials. The networks made a single error after each rule switch, and were able to quickly switch to the new rule and maintain good performance (Figure 1c, d). Correspondingly, single neurons from both modules exhibited rule-modulated persistent activity that lasted several trials (Supplementary Figure 1).

Our networks can reliably maintain good performance after a single correct trial in the new rule, which matches the behavior of monkeys in some previous studies (e.g. Figure 1e). However, the performance of monkeys during this task showed substantial variability across different studies as well as different sessions with a study. The number of error trials that monkeys take to switch to the new rule ranges from one trial to tens of trials [23, 10, 11, 8]. One reason is that performance typically reaches a certain criterion (e.g. 80% correct) but not 100% before rule switching, therefore an error signal could mean an erroneous sensori-motor transformation rather than rule change. Indeed, when training of our model was stopped at 80% rather than 100% accuracy, the resulting network showed gradual switching (Figure 1f, Methods). This point will be addressed further in the Discussion section. In the following sections, we will “open the black box” to understand the mechanism the networks used to perform the WCST.

### Two rule attractor states in the PFC module maintained by interactions between modules

We first dissected the PFC module, which was trained to represent the rule. Since there are two rules in the WCST task we used, we expected that the PFC module might have two attractor states corresponding to the two rules. Therefore, we examined the attractor structure in the dynamical landscape of the PFC module by initializing the network at states chosen randomly from the trial, and evolving the network autonomously (without any input) for 500 time steps (which equals 5 seconds in real time). Then, the dynamics of the PFC module during this evolution was visualized by applying principal component analysis to the population activity. The PFC population activity settled into one of two different attractor states depending on the rule that the initial state belongs to (Supplementary Figure 2a). Therefore, there are two attractors in the dynamical landscape of the PFC module, corresponding to the two rules.

Historically, persistent neural activity corresponding to attractor states were first discovered in the PFC [29, 30, 31, 32]. However, more recent experiments found persistent neural activity in multiple brain regions, suggesting that long-range connections between brain regions may be essential for generating persistent activity [33, 34, 35, 36]. Inspired by these findings, we wondered if the PFC module in our network could support the two rule attractor states by itself, or that the long-range connections between the PFC and the sensorimotor module are necessary to support them. To this end, we lesioned the inter-modular connections in the trained networks and repeated the simulation. Interestingly, we found that for the majority of the trained networks (42 out of 52 for the fast switching networks and 28 out of 28 for the slow switching networks), their PFC activity settled into a trivial fixed point corresponding to an inactive state (Supplementary Figure 2b, c). This result shows that the two rule attractor states in these networks are dependent on the interactions between the PFC and the sensorimotor modules, although a significant part of the excitation still comes from local populations (Supplementary Figure 2d)

### Two emergent subpopulations of excitatory neurons in the PFC module

For the PFC module to keep track of the currently valid rule, the module should stay in the same rule attractor state after receiving positive feedback, but transition to the other rule attractor state after receiving negative feedback. We reasoned that this network function might be mediated by single neurons that are modulated by the task rule and negative feedback, respectively. Therefore, we set out to look for these single neurons.

In the PFC module of the trained networks, there are indeed neurons whose activity is modulated by the task rule in a sustained fashion (example neurons in Supplementary Figure 1 and Figure 2a, top). In contrast, there are also neurons that show transient activity only after negative feedback. Furthermore, this activity is also rule-dependent. In other words, their activity is conjunctively modulated by negative feedback and the task rule (example neurons in Supplementary Figure 1, red traces and Figure 2a, bottom). We termed these two classes of neurons “rule neurons” and “conjunctive error x rule neurons” respectively.

**Figure 2:**
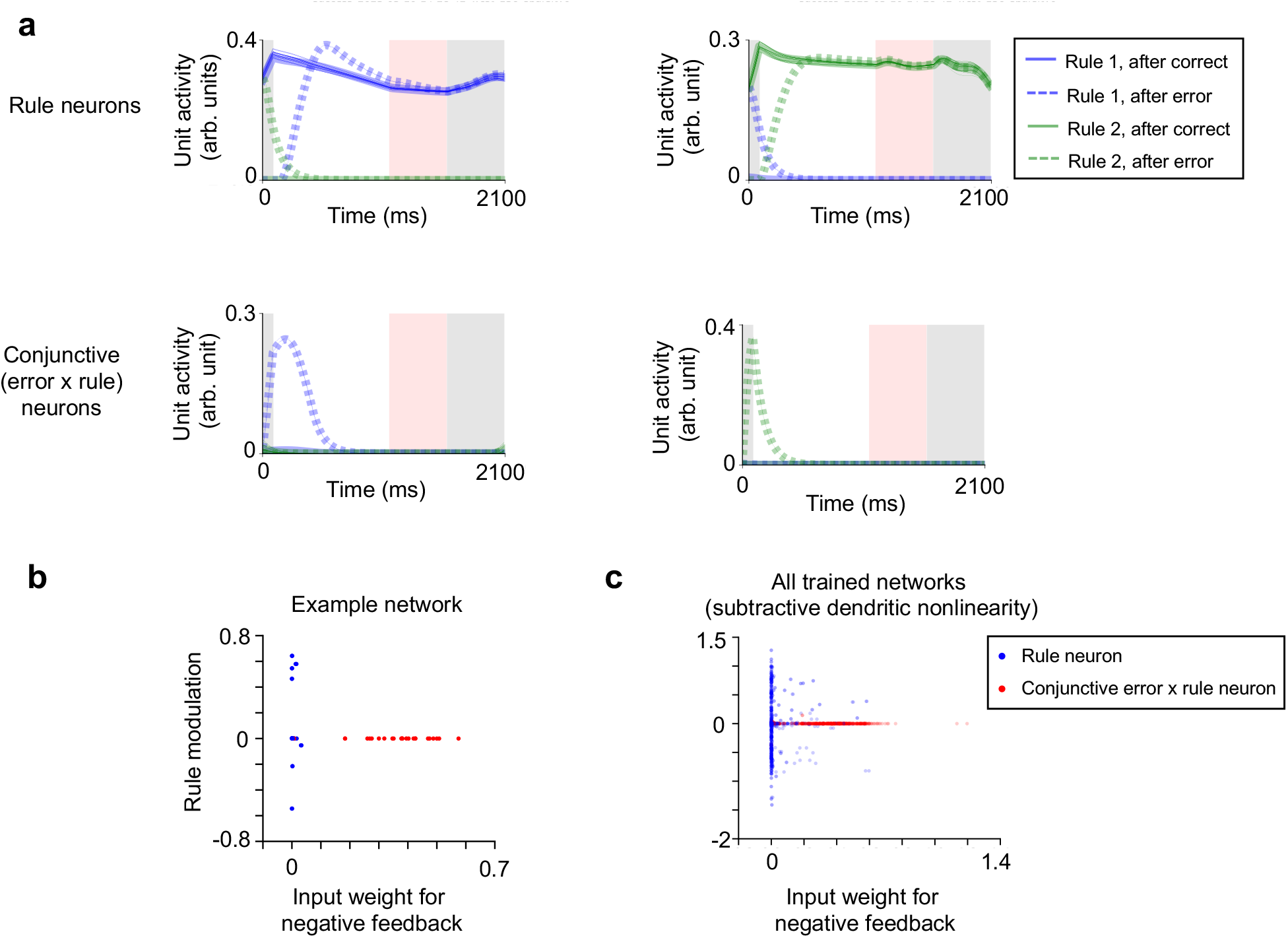
Emergence of two subpopulations of excitatory neurons in the PFC module after training. **a**. Two example rule neurons (top) and conjunctive error x rule neurons (bottom). The solid traces represent the mean activity across trials that follow a correct trial, when those trials belong to color rule (blue) or shape rule (green) blocks. The dashed traces represent the mean activity after error trials, when those trials belong to color rule (blue) or shape rule (green) blocks. We use rule 1 and color rule, as well as rule 2 and shape rule interchangeably hereafter. **b**. Rule neurons and conjunctive neurons are separable. The *x* axis represents the input weight for negative feedback, and the *y* axis is the difference between the mean activity over color rule trials and shape rule trials (for trials following a correct trial). As shown, the rule neurons (blue points) receive little input about negative feedback, but their activity is modulated by rule; The conjunctive error x rule neurons (red points) receive substantial input about negative feedback, but their activity is not modulated by rule (during trials following a correct trial). **c**. The trend in **b** is preserved across a collection of trained networks. Here the results are shown for networks with subtractive dendritic nonlinearity. Networks with divisive dendritic nonlinearity show a similar pattern (Supplementary Figure 3).

We identified all the rule neurons and conjunctive error x rule neurons in the PFC module using a single neuron selectivity measure (see Methods for details). The two classes of neurons are clearly separable on the two-dimensional plane in Figure 2c, where the *x* axis is the input weight for negative feedback, and the *y* axis is the rule modulation, which is the difference in the mean activity between the two rules (for trials following a correct trial). As shown in Figure 2c, rule neurons receive negligible input about negative feedback, and many of them have activity modulated by rule. On the other hand, conjunctive error x rule neurons receive a substantial amount of input about negative feedback, yet their activity is minimally modulated by rule on trials following a correct trial (Figure 2b). This pattern was preserved when aggregating across trained networks (Figure 2c and Supplementary Figure 3). Interestingly, neurons with similar tuning profiles have been reported in the PFC and posterior parietal cortex of macaque monkeys performing the same WCST analog [10, 11].

Across different cell types in the PFC module, on average 23.1% of excitatory neurons, 57.3% of PV neurons and 38.1% of SST neurons were classified as rule neurons in each model. Compared to excitatory neurons, a much smaller fraction of inhibitory neurons in the PFC were classified as conjunctive error x rule neurons. On average, 22.9% excitatory neurons were conjunctive error x rule neurons in each model, compared with 11.5% PV neurons and 5.2% SST neurons. Therefore, we focus only on the excitatory conjunctive error x rule neurons in the analysis below.

We also performed the same analysis on the trained networks that switch rules more slowly (e.g. Figure 1f). In those networks there is a weaker separation between the two subpopulations of excitatory neurons in the PFC module (Supplementary Figure 6a).

### Maintaining and switching rule states via structured connectivity patterns between subpopulations of neurons within the PFC module

Given the existence of rule neurons and conjunctive error x rule neurons, what is the connectivity between them that enables the PFC module to stay in the same rule attractor state when receiving correct feedback, and switch to the other rule attractor state when receiving negative feedback?

To this end, we examined the connectivity between different subpopulations of neurons in the PFC module explicitly, by computing the mean connection strength between each pair of subpopulations. This analysis reveals that the excitatory rule neurons and PV rule neurons form a classic winner-take-all network architecture [37] with selective inhibitory populations [38, 39], where excitatory neurons preferring the same rule are more strongly connected, and they also more strongly project to PV neurons preferring the same rule (Figure 3a). On the other hand, PV neurons project more strongly to both excitatory neurons and other PV neurons with the opposite rule preference (Figure 3a). This winner-take-all network motif together with the excitatory drive from the sensorimotor module (Supplementary Figure 2) is able to sustain one of the two attractor states.

**Figure 3:**
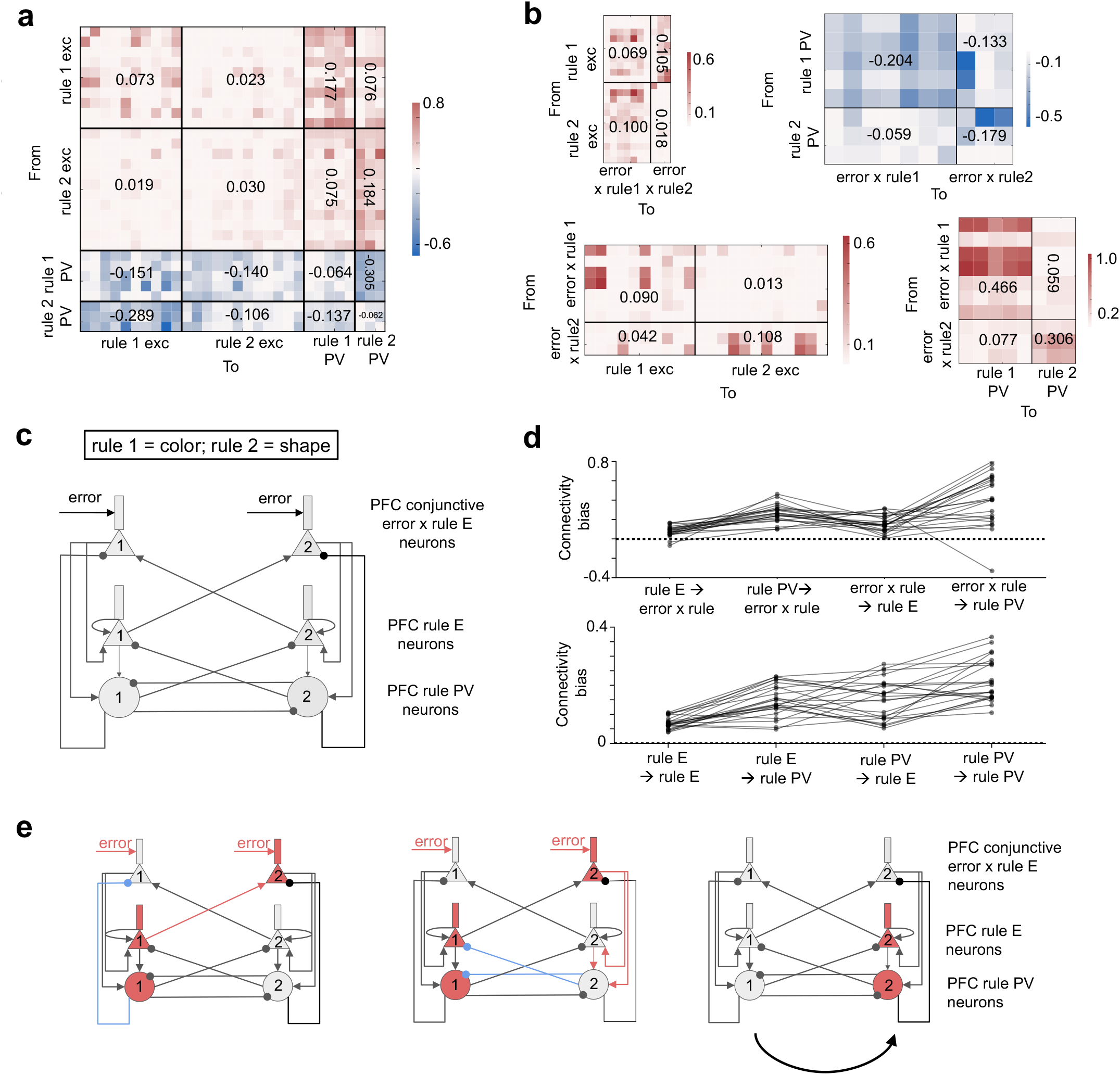
An emergent circuit wiring diagram in the PFC module enables un-cued switching between rule attractor states. **a**. The connectivity matrix between different populations of rule neurons, for an example model. Text indicates the mean connection strength between two populations. Exc: excitatory neuron. **b**. The connectivity between rule neurons and conjunctive error x rule neurons, for an example model. Top left: excitatory rule neurons project more strongly to the conjunctive error x rule neurons that prefer the opposite rule; Top right: PV rule neurons project more strongly to conjunctive error x rule neurons that prefer the same rule; Bottom left: conjunctive error x rule neurons project more strongly to the excitatory rule neurons that prefer the same rule; Bottom right: conjunctive error x rule neurons project more strongly to the PV rule neurons that prefer the same rule. Exc: excitatory neuron. **c**. The simplified circuit diagram between rule neurons and conjunctive neurons based on the result of **b**. The weaker connections are not drawn. Rule 1 represents the color rule and rule 2 represents the shape rule. E: excitatory. **d**. The connectivity biases across all trained models are mostly above 0, both for the connection among rule neurons (top) and the connection between rule neurons and conjunctive error x rule neurons (bottom). Here the results are shown for all networks with subtractive dendritic nonlinearity. Networks with divisive dendritic nonlinearity show similar result (Supplementary Figure 4). E: excitatory neuron. **e**. A schematic showing how the simplified circuit can switch from the rule 1 attractor state to the rule 2 attractor state after receiving the input about negative feedback. When the network is under rule 1 and negative feedback is received, the convergent excitation onto the error x rule 2 neurons (red arrows) and the inhibition onto error x rule 1 neurons (blue arrows) make error x rule 2 neurons more active (left panel). They then excite the rule 2 excitatory and PV neurons, which inhibit the currently-active rule 1 populations (middle panel). As a result, the rule 2 populations become more active than the rule 1 populations and the network switches to rule 2 (right panel). E: excitatory.

Next, how are the rule neurons connected with the conjunctive error x rule neurons such that the sub-network formed by the rule neurons can switch from one attractor to the other in the presence of the negative feedback input? Using the same method, we found that the connectivity between the rule neurons and the conjunctive error x rule neurons exhibited an interesting structure: the excitatory rule neurons more strongly target the conjunctive error x rule neurons that prefer the opposite rule; the PV rule neurons more strongly target conjunctive error x rule neurons that prefer the same rule (Figure 3b, top two panels). On the other hand, the conjunctive error x rule neurons more strongly target the excitatory and PV rule neurons that prefer the same rule (Figure 3b, bottom two panels).

This connectivity structure gives rise to a simple circuit diagram of the PFC module (Figure 3c), which leads to an intuitive explanation of the circuit mechanism underlying the switching of rule attractor state. For example, suppose the network is in the attractor state corresponding to color rule, and has just received a negative feedback and is about to switch to the attractor corresponding to the shape rule (Figure 3e, left). As shown in Figure 2b-c, the input current that represents the negative feedback mainly targets the conjunctive error x rule neurons. In addition, since the network is in the color rule state, the excitatory and PV neurons that prefer the color rule are more active than those that prefer the shape rule. According to Figure 3b (top two panels), the excitatory neurons that prefer the color rule strongly excite the error x shape rule neurons, and the PV neurons that prefer the color rule strongly inhibit the error x color rule neurons. Therefore, the error x shape rule neurons receive stronger total excitation than the error x color rule neurons, and will be more active (Figure 3e, middle). Their activation will in turn excite the excitatory neurons and PV neurons that prefer the shape rule (Figure 3b, bottom two panels). Finally, due to the winner-take-all connectivity between the rule populations (Figure 3a), the excitatory and PV neurons that prefer the color rule will be suppressed, and the network will transition to the attractor state for the shape rule (Figure 3e, right).

It is worth noting that the same mechanism can also trigger a transition in the opposite direction (from shape rule to color rule) in the presence of the same negative feedback signal. This is enabled by the biased connections between the rule and conjunctive error x rule populations.

Is the simplified circuit diagram (Figure 3c) consistent across trained networks, or different trained networks use different solutions? To examine this question, we computed a “connectivity bias” measure between each pair of populations for each trained network. This measure is greater than zero if the connectivity structure between a pair of populations is closer to the one in the simplified circuit diagram in Figure 3c than to the opposite (see Methods for details). Across trained networks, we found that the connectivity biases were mostly greater than zero (Figure 3d), indicating that the same circuit motif for rule maintenance and switching underlies the PFC module across different trained networks.

A similar circuit architecture exists between the excitatory neurons and the SST neurons in the PFC module (Supplementary Figure 5), where SST neurons receive stronger excitatory input from the conjunctive error x rule neurons that prefer the same rule, and also more strongly inhibit the error x rule neurons that prefer the same rule. In addition, they form a winner-take-all connectivity with the rule excitatory neurons by receiving stronger projections from the rule neurons that prefer the same rule and projecting back more strongly to the rule neurons that prefer the opposite rule. Therefore, they contribute to rule maintenance and switching in a similar way as the PV neurons.

When the same analysis was performed on the slow-switching networks (e.g. Figure 1f), we found that although the rule neurons in these networks also form a winner-take-all connectivity structure, the connectivity between error x rule neurons and rule neurons does not exhibit the same structure as in the fast-switching models (Supplementary Figure 6b, c). Therefore, the slow-switching networks have a similar sub-network that encodes the rule, but a poorly organized sub-network between the error x rule and rule neurons, which may explain why switching the rule takes more trial-and-error in these networks.

### Top-down propagation of the rule information through structured long-range connections

Given that the PFC module can successfully maintain and update the rule representation, how does it use the rule representation to reconfigure the sensorimotor mapping?

First, we found that neurons in the sensorimotor module were tuned to rule (Supplementary Figure 7a), since they receive top-down input from the rule neurons in the PFC module. The PFC module exerts top-down control through three pathways: the monosynaptic pathway from the excitatory neurons in the PFC module to the excitatory neurons in the sensori-motor module, the tri-synaptic pathway that goes through the VIP and SST neurons in the sensorimotor module, and the di-synaptic pathway mediated by the PV neurons in the sensorimotor module (Figure 1b). We found that there are structured connectivity patterns along all three pathways. Along the monosynaptic pathway, excitatory rule neurons in the PFC module preferentially send long-range projections to the excitatory neurons in the sensorimotor module that prefer the same rule (Figure 4a). Along the tri-synaptic pathway, PFC excitatory rule neurons also send long-range projections to the SST and VIP interneurons in the sensorimotor module that prefer the same rule (Figure 4b-c). The SST neurons in turn send stronger inhibitory connections to the dendrite of the local excitatory neurons that prefer the opposite rule (Figure 4d). Along the di-synaptic pathway, the PV neurons are also more strongly targeted by PFC excitatory rule neurons that prefer the same rule (Figure 4e), and they inhibit more strongly the local excitatory neurons that prefer the opposite rule (Figure 4f). These trends are mostly preserved across trained networks (Supplementary Figure 8). Therefore, rule information is communicated to the sensorimotor module synergistically via the mono-synaptic excitatory pathway, the tri-synaptic pathway that involves the SST and VIP neurons, as well as the di-synaptic pathway that involves the PV neurons, as illustrated in Figure 4g.

**Figure 4:**
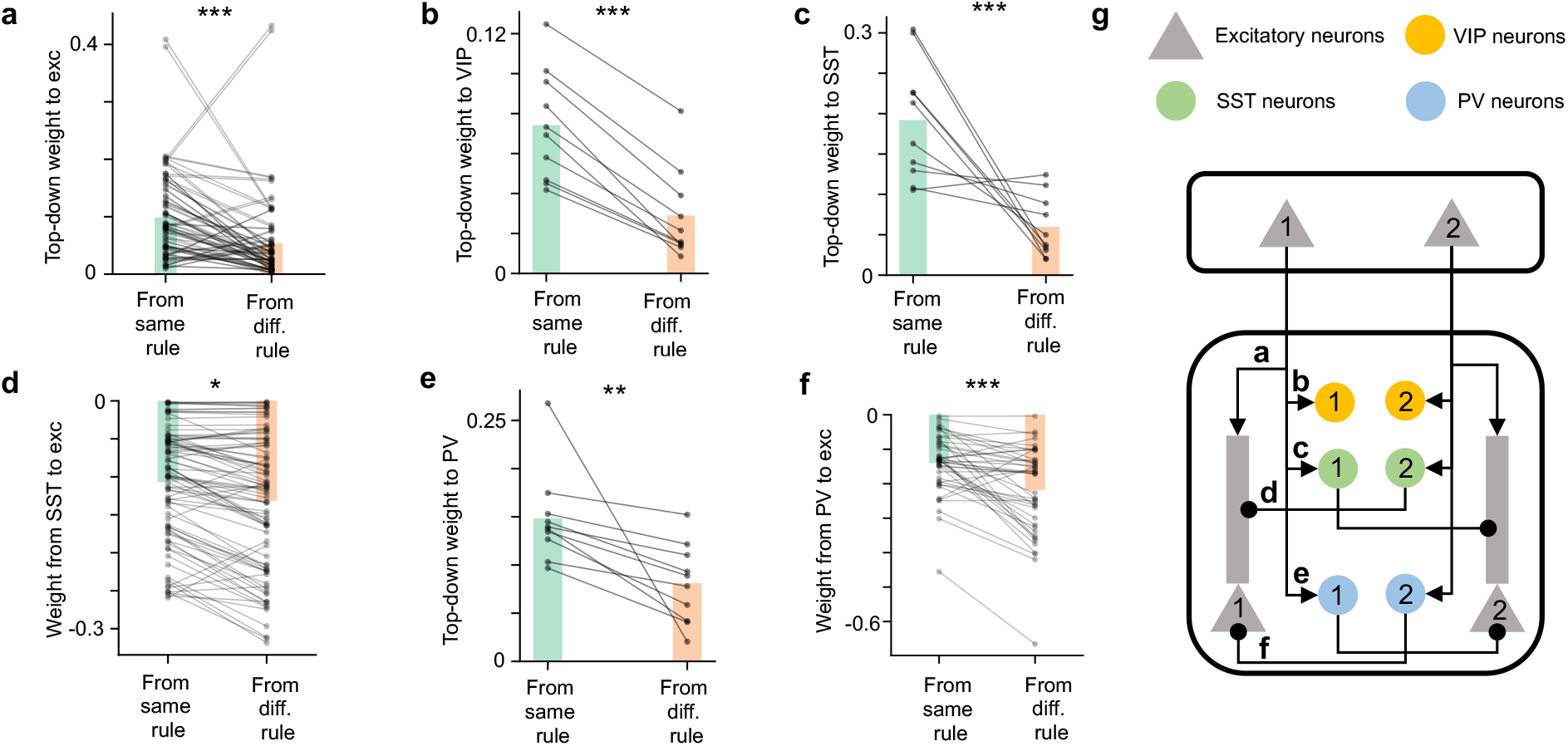
Structured top-down connections enable the propagation of the rule information. **a**. Each line represents the mean connection strength onto one excitatory neuron in the sensorimotor module, from the PFC excitatory neurons that prefer the same rule and the opposite rule. Bars represent mean across neurons. PFC excitatory neurons project more strongly to excitatory neurons in the sensorimotor module that prefer the same rule (one-sided Student’s t test, *p* = 3.0 ∗ 10^−5^, *n* = 88 dendritic branches. Only target neurons with a non-zero rule selectivity were included). Exc: excitatory neurons. **b**. The same data but for the connections from PFC excitatory neurons to VIP neurons in the sensorimotor module (one-sided Student’s t test, *p* = 4.1 ∗ 10^−4^, *n* = 10 VIP neurons). **c**. The same data but for the connections from PFC excitatory neurons to SST neurons in the sensorimotor module (one-sided Student’s t test, *p* = 4.3 ∗ 10^−5^, *n* = 10 SST neurons). **d**. The same data but for the connections from SST neurons to the excitatory neurons in the sensorimotor module (one-sided Student’s t test, *p* = 0.02, *n* = 88 dendritic branches. Only target neurons with a non-zero rule selectivity were included). Exc: excitatory neurons. **e**. The same data but for the connections from PFC excitatory neurons to PV neurons in the sensorimotor module (one-sided Student’s t test, *p* = 1.6 ∗ 10^−3^, *n* = 10 PV neurons. **f**. The same data but for the connections from PV neurons to excitatory neurons in the sensorimotor module (one-sided Student’s t test, *p* = 6.3∗ 10^−4^, *n* = 44 excitatory neurons). Exc: excitatory neurons. **g**. The structure of the top-down connections as indicated by the results in **a**-**f**. The weaker connections are not shown. Results in **a**-**f** are shown for an example network with subtractive dendritic nonlinearity. Networks with divisive and subtractive dendritic nonlinearity show similar patterns (Supplementary Figure 8).

### Structured input and output connections of the sensorimotor module enable rule-dependent action selection

Given the top-down rule information from the PFC module, how does the sensori-motor module implement the sensorimotor transformation (from the cards to the response to one of the three spatial locations)? We sought to identify the structures in the input, recurrent and output connections of the sensorimotor module that give rise to this function.

We started by observing that excitatory neurons in the sensorimotor module showed a continuum of encoding strengths for task rule, response location and card features, and many neurons show conjunctive selectivity for these variables (Figure 5b, Supplementary Figure 7b). Therefore, we assigned each excitatory neuron in the sensorimotor module a preferred rule *R*, a preferred response location *L* and a preferred shared feature *F*, which is the feature that is present in both the reference card and the test card at *L*. For example, neurons with *R* = *color rule, L* = 1 and *F* = *blue* have the highest activity during color rule trials when the correct response is to choose the test card at location 1, and when that test card shares the blue color with the reference card (it belongs to the population with the filled green color in Figure 5a). Intuitively, for this group of neurons to show such selectivity, they should receive strong input from the input neurons that encode the *F* = *blue* feature of the test card at location *L* = 1 and the same feature for the reference card. This would enable them to detect when the test card at *L* = 1 and the reference card both have the feature *F* = *blue*.

**Figure 5:**
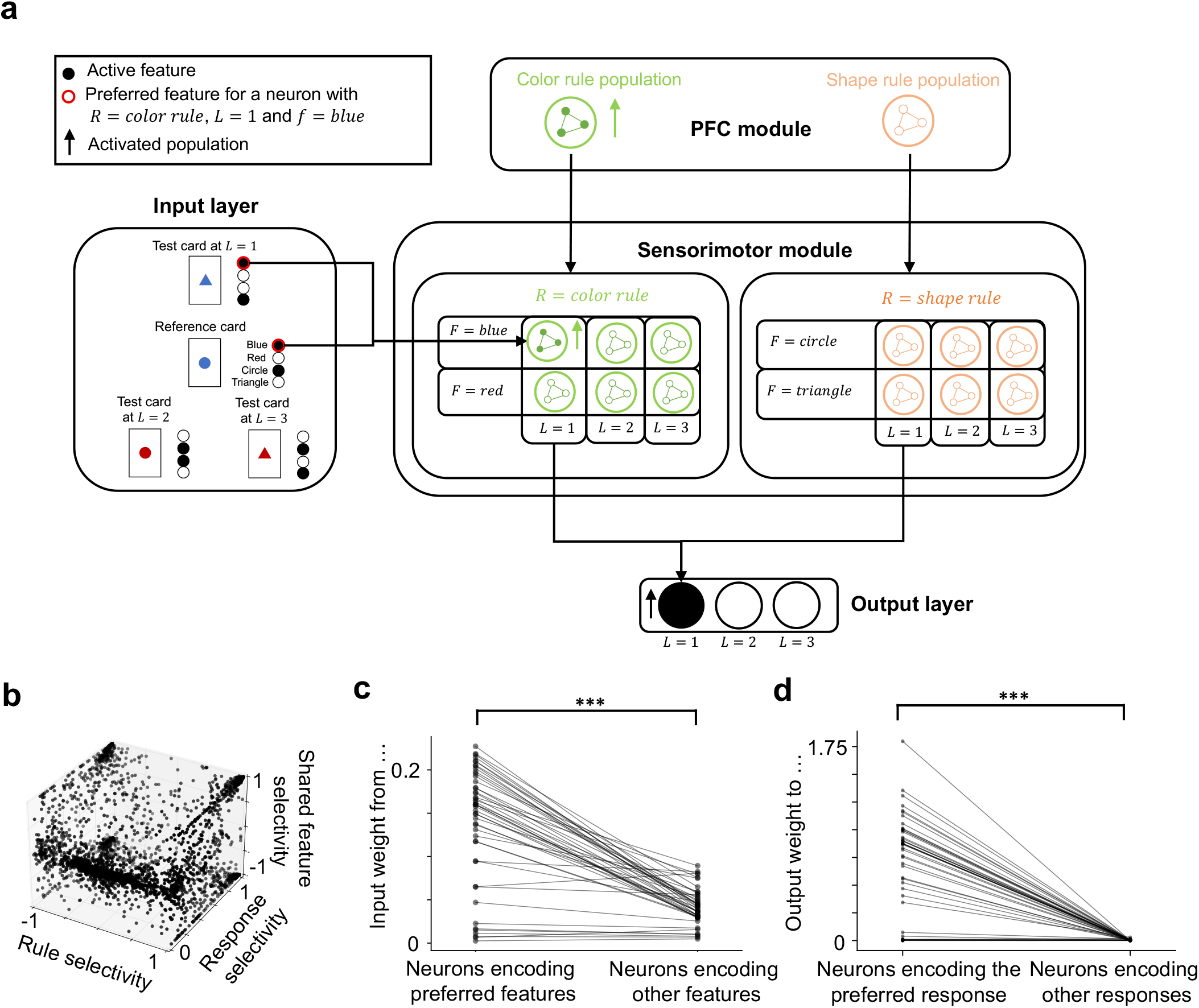
Structures in the input and output weights of the sensorimotor module enable rule-dependent action selection. **a**. Excitatory neurons in the sensorimotor module were classified according to their preferred rule *R*, response location *L* and shared feature *F*. For example, neurons with *R* = *color rule, L* = 1 and *F* = *blue* have the highest activity during color rule trials when the network chooses the test card at *L* = 1, and when that card shares the *F* = *blue* feature with the reference card. For a neuron with a given *R, L* and *F*, its “preferred features” are defined as the feature *F* of the reference card and same feature of the test card at location *L*. For example, the preferred features for the above neurons with *R* = *color rule, L* = 1 and *F* = *blue* are the blue feature of the reference card and the test card at *L* = 1. **b**. The joint distribution of the selectivity for rule (*R*), response location (*L*) and shared feature (*F*) across all neurons in the sensorimotor module. Result is aggregated across all trained networks. **c**. Excitatory neurons in the sensorimotor module receive stronger connections from the input neurons that encode their preferred features (as defined in **a**). Each line represents one excitatory neuron in the sensorimotor module. One-sided Student’s t-test, *p* = 2.4 ∗ 10^−15^, *n* = 47 neurons. **d**. Excitatory neurons in the sensorimotor module send stronger connections to the output neuron that represents their preferred response location. Each line represents one excitatory neuron in the sensorimotor module. One-sided Student’s t-test, *p* = 2.3∗ 10^−17^, *n* = 45 neurons.

In general, for neurons that prefer rule *R*, response location *L* and shared feature *F*, we can define their “preferred features” as the feature *F* of the reference card and the same feature *F* for the test card at location *L*. Across all excitatory neurons in the sensorimotor module, we found that the connections from the input neurons that encode these preferred features were significantly stronger than the connections from the input neurons that encode other features (Figure 5c). In addition, there is also an intuitive structure in the output connections, where excitatory neurons in the sensorimotor module that prefer a particular response location send stronger connections to the output neuron that represents the same response location (Figure 5d). These patterns were found to be consistent across trained networks (Supplementary Figure 9).

These structures in the input and output connections give rise to an intuitive explanation of how the sensorimotor module can perform rule-dependent action selection required for the WCST. Here we illustrate this mechanism with an example trial (Figure 5a), where the current rule is color, the reference card is a blue circle, and the test cards at locations 1 2 and 3 are blue triangle, red circle and red triangle, respectively. According to the rule of WCST, the correct response should be location 1, since the test card at that location matches the reference card in color. This choice can be generated as follows: the excitatory population in the sensorimotor module that prefers *R* = *color rule, L* = 1 and *F* = *blue* will be most strongly activated since they not only receive strong top-down excitatory input from the PFC module, but also the strongest excitatory input from the input neurons. Therefore, they are the most strongly activated population (Figure 5a). Since they prefer response location 1, they will activate the output neuron that prefers response location *L* = 1, which is the correct response.

Panel **a** shows an example trial that illustrates how the sensorimotor module can generate the correct response. See text for the detailed mechanism.

### Recurrent connectivity and dynamics within the sensorimotor module

Given that different populations of neurons in the sensorimotor module receive differential inputs about the external sensory stimuli and rule via the structured input and top-down connections, how are they recurrently connected to produce dynamics that lead to a categorical choice? To answer this, we first visualized the population neural dynamics in the sensorimotor module by using principal component analysis (Figure 6a-b). As shown in Figure 6a, neural trajectories during the inter-trial interval are clustered according to the task rule. During the response period, the neural trajectories are separable according to the response locations, albeit only in higher-order principal components (Figure 6b). In addition, the subspaces spanned by neural trajectories of different rules and response locations are more orthogonal to each other compared to randomly shuffled data (Figure 6c-d, Methods).

**Figure 6:**
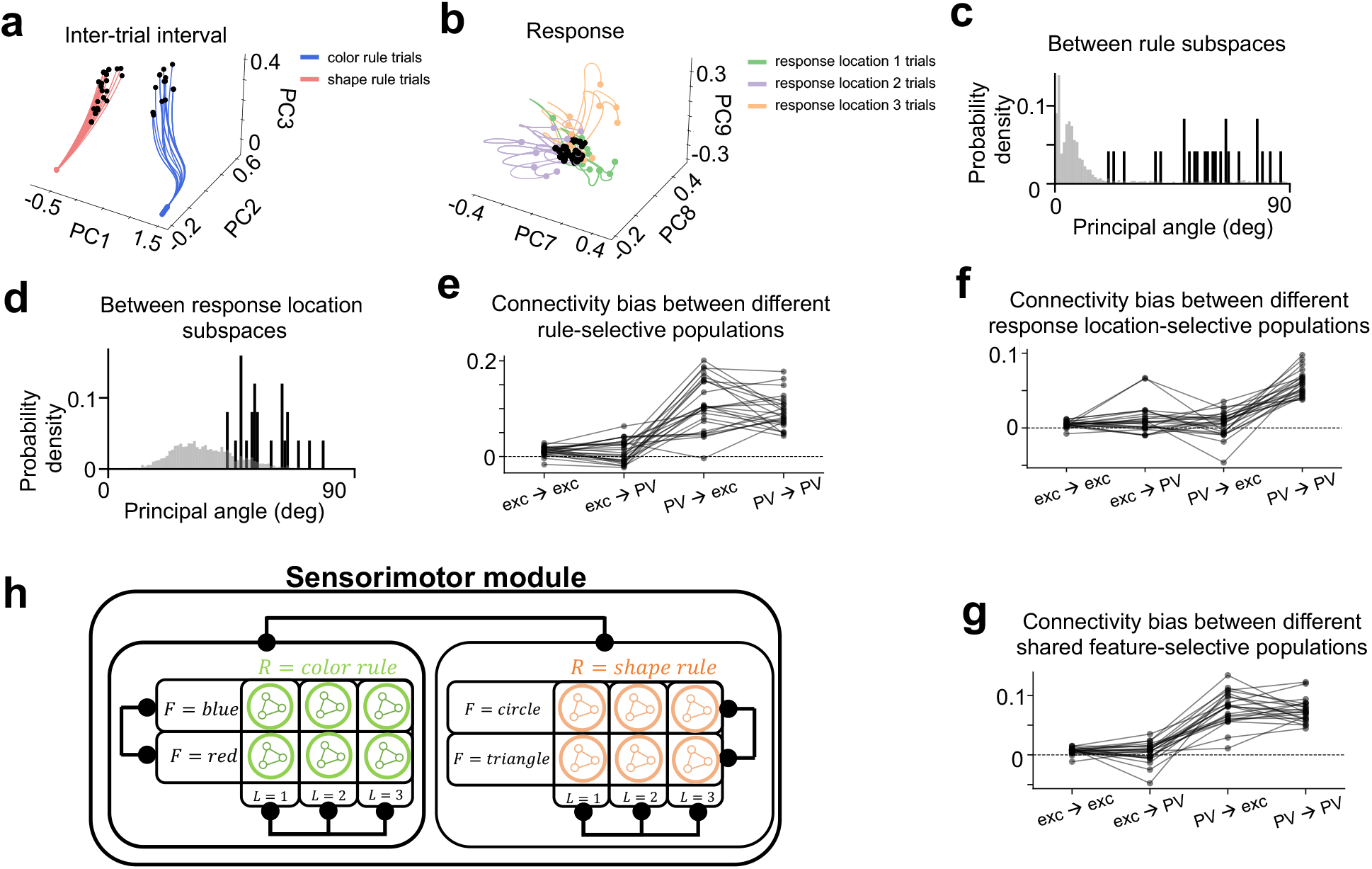
Recurrent dynamics and connectivity within the sensorimotor module. **a**. Neural trajectories during the intertrial interval (ITI) colored by rule, visualized in the space spanned by the first three principal components. Black dots represent the start of the ITI. Only trials following a correct trial were included. **b**. Neural trajectories during the response period colored by response location, visualized in the space spanned by higher order principal components. Black dots represent the start of the response period. Only trials following a correct trial were included. **c**. The principal angle between the subspaces spanned by neural trajectories during different task rules (gray distribution represents the principal angle obtained through shuffled data, see Methods). Each data point represents one trained network. **d**. The principal angle between the subspaces spanned by neural trajectories during different responses (gray distribution represents the principal angle obtained through shuffled data, see Methods). Each data point represents one trained network. **e**. The connectivity biases between different rule-selective populations across models. Exc: excitatory neurons. **f**. The same as **e** but for different response location-selective populations. Exc: excitatory neurons. **g**. The same as **e** but for different shared feature-selective populations. Exc: excitatory neurons. **h**. The results in **e** - **g** show that neural populations selective for different rules, response locations and shared features mutually inhibit each other.Data in **c**-**g** are shown for networks with subtractive dendritic nonlinearity. Networks with divisive dendritic nonlinearity show similar result (Supplementary Figure 10).

What connectivity structure gives rise to this signature in the population dynamics? To answer this, we examined the pattern of connection weights between excitatory and PV neurons that prefer different rules (*R*), response locations (*L*), and shared features (*F*) by computing the connectivity biases between populations of neurons that are selective to different rules (Figure 6e), response locations (Figure 6f) and shared features (Figure 6g). A greater than zero connectivity bias means populations that prefer different rules (or response locations or shared features) form a winner-take-all circuit structure analogous to the one observed between rule-selective populations in the PFC module (c.f. top panel of Figure 3d, details about how the connectivity biases were computed is described in Methods). We observed that many of the connectivity biases were significantly above zero (Figure 6e-g), especially for the ones that correspond to the inhibitory connections originating from the PV neurons. This indicates that populations of neurons in the sensorimotor module that are selective to different rules, response locations and shared features overall inhibit each other. This mutual inhibition circuit structure magnifies the difference in the amount of long-range inputs that different populations receive (Figure 5) and leads to a categorical choice.

### SST neurons are essential to dendritic top-down gating

The previous sections elucidate the key connectivity structures that enable the network to perform the WCST. In this final section we are going to take advantage of the biological realism of the trained RNN and examine the function of SST neurons in this model.

It has been observed that different dendritic branches of the same neuron can be tuned to different task variables [40, 41, 42, 43]. This property may enable individual dendritic branches to control the flow of information into the local network [19, 22]. Given these previous findings, we examined the coding of the top-down rule information at the level of individual dendritic branches. Since each excitatory neuron in our networks is modeled with two dendritic compartments, we examined the encoding of rule information by different dendritic branches of the same excitatory neuron in the sensorimotor module.

One strategy of gating is for different dendritic branches of the same neuron to prefer the same rule, in which case these neurons form distinct populations that are preferentially recruited under different task rules (population-level gating, Figure 7a, right). An alternative strategy is for different dendritic branches of the same neuron to prefer different rules, which would enable these neurons to be involved in both task rules (dendritic branch-specific gating, Figure 7a, left).

**Figure 7:**
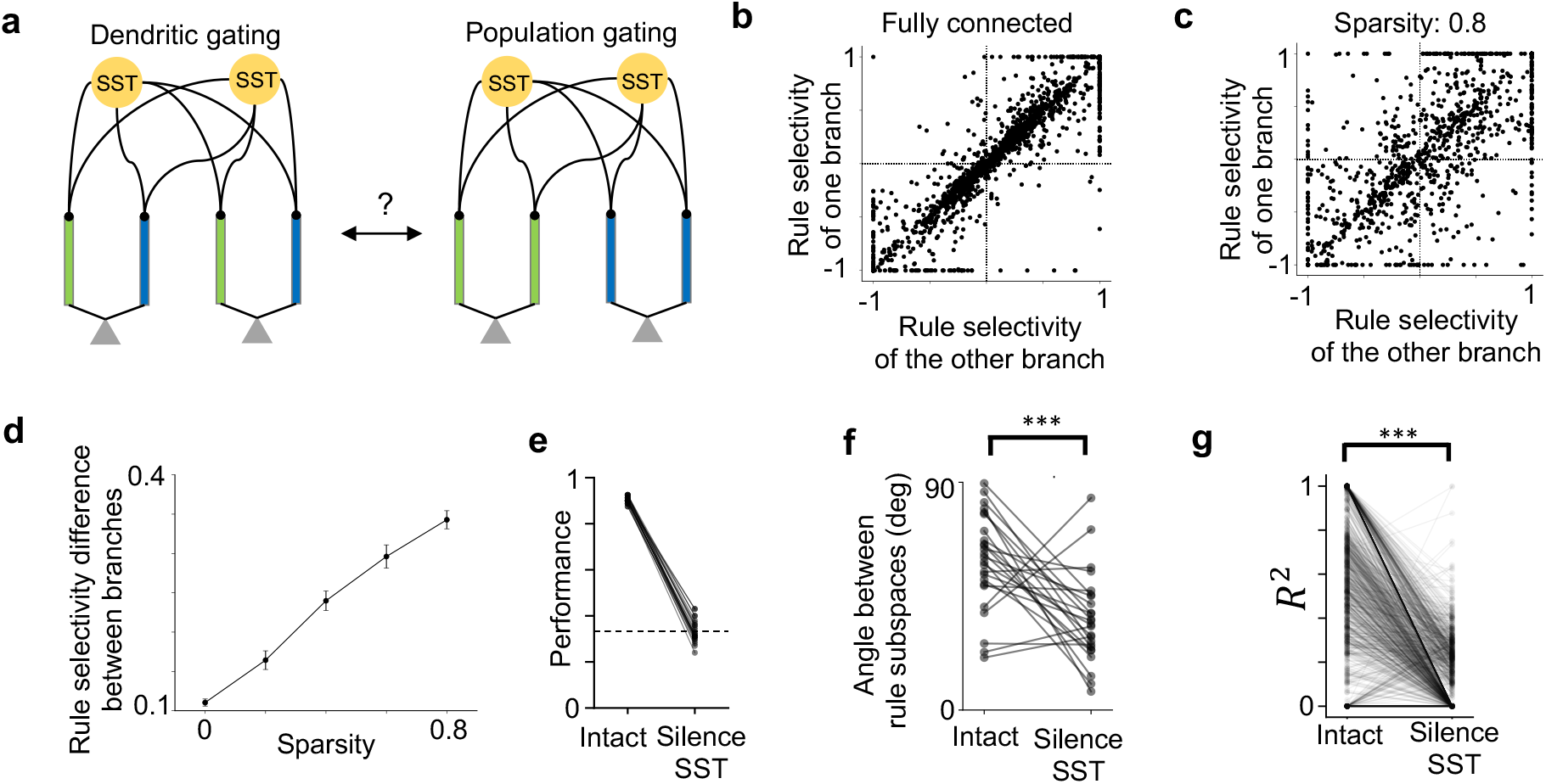
Examining the role of SST neurons in the sensorimotor module in top-down gating. **a**. Two scenarios for top-down gating. Blue and green color represent dendritic branches that prefer one of the two rules. Different dendritic branches of the same neuron could have similar (right) or different (left) rule selectivity. **b**. The rule selectivity of one dendritic branch against the other, aggregated across all models where the connections from the SST neurons to the excitatory neurons are all-to-all. The rule selectivity for different dendritic branches of the same neuron are highly correlated. **c**. The rule selectivity of one dendritic branch against the other, aggregated across all models where the 80% of the connections from the SST neurons to the excitatory neurons were frozen at 0 throughout training. Note the the rule selectivity for different dendritic branches of the same neuron are less correlated than in **b**. **d**. The degree of dendritic branch-specific encoding of the task rule is quantified as the difference in the rule selectivity between the two dendritic branches of the same excitatory neuron in the sensorimotor module. Across all dendritic branches, this quantity increases with the sparsity of the SST→ dendrite connectivity. Error bars represent standard error of the mean across *n* = 1890, 350, 490, 560, 980 pairs of dendritic branches for sparsity levels 0, 0.2, 0.4, 0.6, 0.8 respectively. **e**. Task performance drops significantly after silencing SST neurons in the sensorimotor module. Each line represents a trained network. **f**. The principal angle between rule subspaces (c.f. Figure 6c) drops significantly after silencing SST neurons in the sensorimotor module (One-sided Student’s t-test, *p* = 2.7 ∗ 10^−5^, *n* = 25 networks). Each line represents a trained network. **g**. The strength of conjunctive coding of rule and stimulus (as measured by the *R*^2^ value in a linear model with conjunctive terms, see Methods) decreases after silencing SST neurons in the sensorimotor module (One-sided Student’s t-test, *p* = 1.7 ∗ 10^−175^, *n* = 1750 neurons). Each line represents one neuron. Results were aggregated across networks.

In light of this, we examined for our trained networks to what extent they adopt these strategies. We found that the rule selectivity between different dendritic branches of the same neuron were highly correlated (Figure 7b). This indicates that the trained networks are mostly using the population-level gating strategy, where different dendritic branches of the same neuron encode the same rule.

What factors might determine the extent to which the trained networks adopt these two strategies? Previous modeling work suggests that sparse connectivity from SST neurons to the dendrites of the excitatory neurons increases the degree of dendritic branch-specific gating, in the case where the connectivity is random (Figure 4f in Ref.[22]). To see if the same effect is present in our task-optimized network with structured connectivity, we retrained networks with different levels of sparsity from 0 to 0.8 and studied its effect on the dendritic branch specificity of rule coding (Methods). We found that the degree of dendritic branch-specific encoding of the task rule increased with sparsity (see Figure 7c, d for subtractive dendritic nonlinearity; Supplementary Figure 11a for divisive dendritic nonlinearity). Intuitively, when the connection is sparse, a smaller number of SST neurons target each dendritic branch, making it more likely that the branch receives an uneven number of inputs from SST neurons selective for different rules. Taken together, we observed that the trained networks adopted a mixture of population-level and dendritic-level gating strategies for top-down control, and the balance between the two strategies depended on the sparsity of the connections from the SST neurons to the dendrites of excitatory neurons.

Indeed, SST neurons play a causal role in relaying the top-down rule information into the sensorimotor network and reconfiguring its dynamics according to the task rule. We simulated optogenetic inhibition by silencing the SST neurons in the sensorimotor module, which significantly impaired task performance (Figure 7e, see Methods section for details of the protocol). In addition, the principal angle between the subspaces for different rules (Figure 6c) significantly decreased after SST neurons in the sensorimotor module were silenced for networks with subtractive dendritic nonlinearity (Figure 7f). This effect was not significant for networks with divisive dendritic nonlinearity (Supplementary Figure 11c). Silencing of the SST neurons in the sensorimotor module also significantly diminished non-linear mixed-selective coding of rule and stimulus among the excitatory neurons in the sensorimotor module (Figure 7g, Supplementary Figure 12, Methods), which has been proposed to be important for rule-based sensorimotor associations [44, 45, 46]. Taken together, these results highlight the role that SST neurons in the sensorimotor module play during top-down control. This analysis also shows that by combining artificial neural network with knowledge from neurobiology, it is possible to probe the functions of fine-scale biological components in cognitive behaviors.

## Discussion

In this paper, we analyzed recurrent neural networks trained to perform a classic task involving un-cued task switching - the Wisconsin Card Sorting Test. The networks consist of a “PFC” module trained to represent the rule and interacts with a “sensorimotor” module that instantiates different sensorimotor mappings depending on the rule. In order to study the functions of dendritic computation and different neuronal types, each module is endowed with excitatory neurons with two dendritic branches as well as three major types of inhibitory neurons - PV, SST and VIP. After training, we dissected the trained networks to elucidate a number of intra-areal and inter-areal neural circuit mechanisms underlying WCST, as summarized in Figure 8.

**Figure 8:**
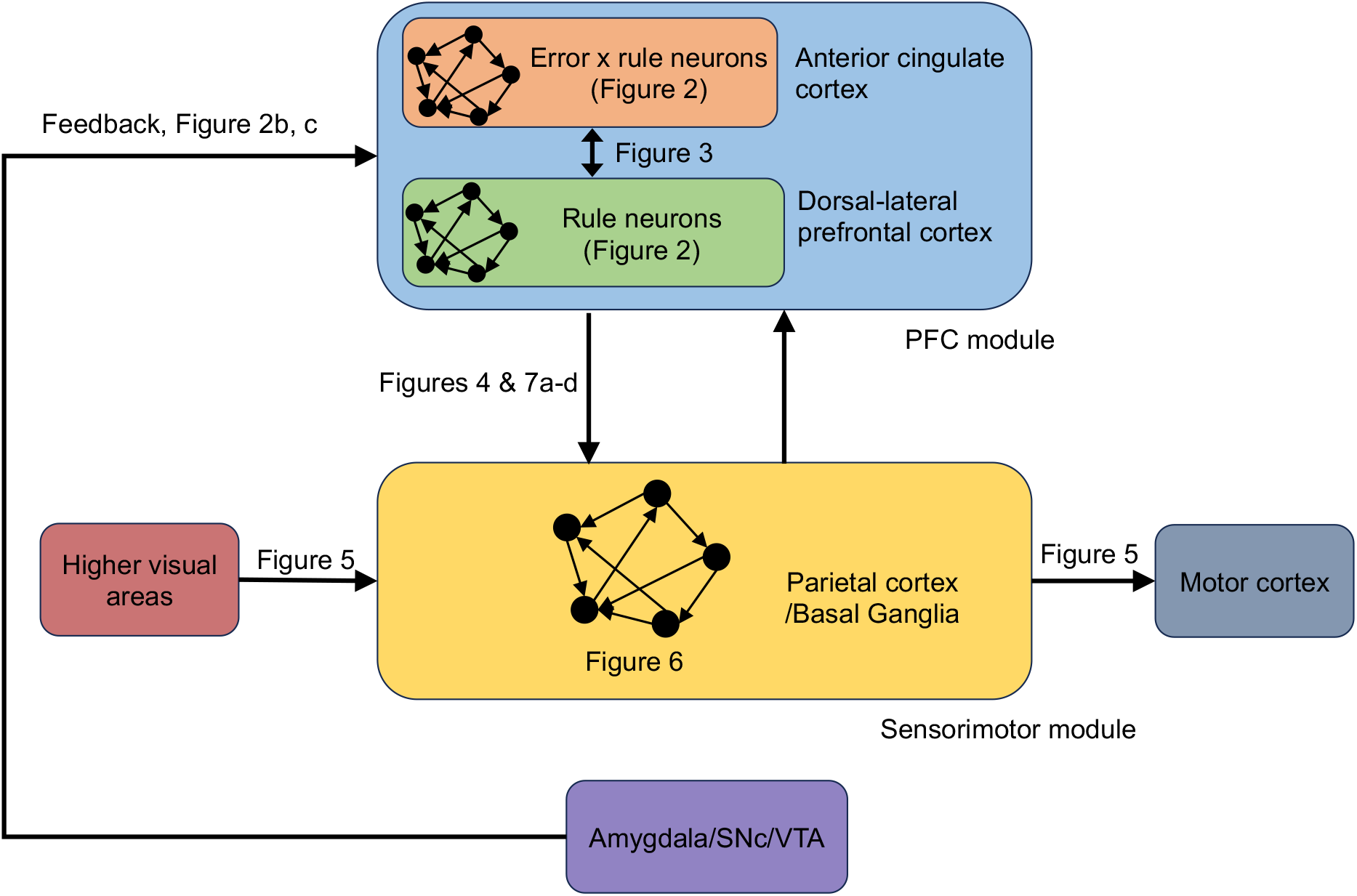
A summary of the main results. Different components of the model can be mapped to different brain regions; The conjunctive error x rule neurons may reside in the anterior cingulate cortex; The rule neurons may be found in the dorsal-lateral PFC; The input to the PFC module about negative feedback may come from subcortical areas such as the amygdala or the midbrain dopamine neurons; The sensorimotor module may correspond to parietal cortex or basal ganglia which have been shown to be involved in sensorimotor transformations; Neurons in the input layer that encode the color and shape of the card stimuli exist in higher visual areas such as the inferotemporal cortex; Neurons in the output layer that encode different response locations could correspond to neurons in the motor cortex.

### Mapping between model components and brain regions

Different components of the trained network can be mapped to different brain regions (Figure 8). While single neurons in the dorsal-lateral PFC (DLPFC) are shown to encode the task rule [47], neurons in the anterior cingulate cortex (ACC) are thought to be important for performance monitoring [48], and have been shown to receive more input about the feedback [49, 50, 51, 52]. Therefore, the rule neurons and conjunctive error x rule neurons in the model correspond to the putative functions of the neurons in DLPFC and ACC. The input to the PFC module about negative feedback may come from subcortical areas such as the amygdala [53] or from the dopamine neurons in the substantia nigra pars compacta (SNc) and ventral tegmental area (VTA) [54, 55]. The sensorimotor module may correspond to parietal cortex or basal ganglia which have been shown to be involved in sensorimotor transformations [56, 57]. The neurons in the input layer that encode the color and shape of the card stimuli exist in higher visual areas such as the inferotemporal cortex [58, 59, 60]. The neurons in the output layer that encode different response locations could correspond to movement location-specific neurons in the motor cortex [61].

Results in **b**-**g** are for networks with subtractive dendritic nonlinearity. See Supplementary Figure 11 for networks with divisive dendritic nonlinearity.

### Attractor states supported by inter-areal connections

We observed that in many networks, the interaction between the two modules was needed to sustain the two rule attractor states (Supplementary Figure 2b,c), although the majority of the excitatory input to PFC neurons come from local population (Supplementary Figure 2d). Traditionally, it was thought that local interactions within the frontal cortex are sufficient for the maintenance of the persistent activity [29, 30, 31, 32]. Recent large-scale electrophysiological recordings, on the other hand, revealed highly distributed encoding of cognitive variables [33, 35, 62, 63, 64, 65]. In addition, distributed patterns of persistent activity emerge in neural network models of multiple brain regions that are constrained by anatomical and neurophysiological data [66, 67]. Despite the empirical evidence, the functional advantages of this multi-areal encoding scheme remain an open question.

### Circuit mechanism in the frontal-parietal network for rule maintenance and update

We found that the PFC module of the trained networks used a particular mechanism for maintaining and updating the rule that depends on the interaction between two distinct populations of neurons that emerge in the PFC module as a result of training: neurons that only encode the rule, and neurons that conjunctively encode negative feedback and rule (Figure 3c). Neurons that show conjunctive selectivity for rule and negative feedback have been reported in monkey prefrontal and parietal cortices while they perform the same WCST task [10, 11]. Theoretical work suggests that these mixed-selective neurons are essential if the network needs to switch between *different* rule attractor states after receiving the *same* input that signals negative feedback [68]. In addition, this circuit mechanism is consistent across dozens of trained networks with different initializations and dendritic nonlinearities (Figure 3d and Supplementary Figure 4).

This circuit mechanism bears resemblance to a previous circuit model of WCST [69]. In that model, different rules are maintained by different “rule-coding clusters” that have self-excitatory connections and lateral inhibitory connections, similar to the winner-take-all attractor network between the rule neurons in our trained networks. On the other hand, these two models use different mechanisms for rule shifting: while in the Dehaene and Changeux model it is achieved via synaptic desensitization caused by the convergence of two inputs (one that signals the recent activation of the synapse, and another that signals negative feedback), our trained networks use structured connections between rule neurons and a separate population of conjunctive error x rule to shift between rules states (c.f. Figure 3c). Notably, neurons that represent the conjunction of negative feedback and rule have been reported in the frontoparietal cortices of monkeys performing WCST [10, 11].

The simplified circuit for the PFC module in Figure 3c can be applied not only to rule switching, but to the switching between other behavioral states as well. For example, it resembles the head-direction circuit in fruit fly [70], where the offset in the connections between the neurons coding for head direction and those coding for the conjunction of angular velocity and head direction enables the circuit to update the head-direction attractor state using the angular velocity input. In addition, this circuit structure may underlie the transition from staying to switching during patch foraging behavior. Indeed, in a laboratory task mimicking natural foraging for monkeys, it was found that neurons in the anterior cingulate cortex increase their firing rates to a threshold before animals switch to another food resource [71].

### Connecting subspace to circuits

Methods that describe the representation and dynamics on the neuronal population level have gained increasing popularity and generated insights that cannot be discovered using single neuron analysis (e.g. [61, 72]). In the meantime, it would be valuable to connect population-level phenomena to their underlying circuit basis [73]. In our model, we found that silencing of the SST neurons has a specific effect on the population-level representation, namely, it decreased the angle between rule subspaces (Figure 7f). Interestingly, this effect was much more prominent in networks with subtractive dendritic inhibition (Figure 7f) than divisive dendritic inhibition (Supplementary Figure 11c). Activating SST neurons can produce both forms of dendritic inhibition, depending on the exactly protocol used [74, 75, 76]. Future work is required to understand the mechanistic link between different forms of dendritic inhibition and the geometry of neural representation.

We also found that silencing the other types of inhibitory neurons in the model has different effects. Silencing the PV neurons led to runaway activity in many networks (Supplementary Figure 11e). Silencing the VIP neurons, on the other hand, caused no significant changes of the task performance (Supplementary Figure 11f). The lack of effect after silencing the VIP neurons is due to the fact that the VIP neurons were largely inhibited by the SST neurons in the trained networks. It is conceivable that VIP neurons would play a more important role if additional constraints are introduced in training RNN to produce a robust disinhibitory motif as observed experimentally [19, 20, 21, 77, 78, 16]. Future work could study the function of VIP neurons under more realistic connectivity constraints between different cell types (e.g. [79]).

### Cued versus un-cued rule switching

The task our networks were trained on is an example of un-cued rule switching task, where the rule is not explicitly cued on each trial but needs to be inferred from the feedback of previous trials. This differs from cued rule switching, another commonly studied rule-based task paradigm where the task rule is explicitly cued during each trial (e.g. [3, 2]). Compared to cued rule switching, additional mechanisms and neural circuitries responsible for inferring the rule shift and maintaining the relevant rule are needed to perform uncued rule switching (although monkeys could employ alternative strategies that require less cognitive load for rule maintenance, as mentioned in [28]).

This difference in the required neural mechanisms leads to a difference of task performance between cued and un-cued rule switching: while it is possible to shift rules with few errors in cued rule switching tasks, such behavior is highly demanding in un-cued rule switching tasks [10, 8]. Below we will further discuss the limitation of our work in terms of the behavior during rule switching.

### Dynamics of behavior during un-cued rule switching

The fast-switching networks in this study switch rules in just one trial (Figure 1c, d). This fast switching agrees with the monkey behavior in some studies [23, 11, 28], but other studies report that monkeys switch rules using on average tens of trials [10, 8]. For example, in Ref.[8], monkeys’ performance is at chance after a single error (Figure 3D in [8]), and they gradually use positive feedback to reinforce their behavior according to the new rule (Figure 4A in [8]). When our network model was trained to achieve less than perfect accuracy, switching after a rule change now takes a few trials (Figure 1f) similar to behavioral observations of many monkey experiments. In this case, the rule-selective neurons in the PFC module still form a winner-take-all attractor network, but their connectivity pattern with the conjunctive error x rule neurons are not as clear cut (Supplementary Figure 6).

Indeed, in WCST and related rule switching paradigms, subjects’ performance is often not perfect even during trials when the rule is fixed. This is possibly because during training, the rule is switched when the subjects’ performance reaches a certain criterion (e.g. 85% correct in a sequence of 20 trials in Ref.[8]). In that case, negative feedback can due to either a rule switch or the inaccuracy in the sensorimotor transformation (even under the correct rule). Therefore, subjects need to integrate information across several trials to decide whether the rule has actually switched. For example, Purcell and Kiani [80] analyzed the behavior of humans in an environment switching task. The task has a similar structure to the WCST analog used in this study, except that the noise level in the stimuli varies from trial to trial. It was shown that the behavior of subjects can be well described by a Bayesian ideal observer model, where the evidence towards an environment switch is incremented whenever the subjects make an error, and the amount of increase depends on the difficulty of the error trial: the easier the error trial is, the more likely that the environment has switched and the larger the incremental evidence towards an environment switch is.

Aside from the difficulty of the task under a fixed rule, the relationship between the different rules may also play a role in how fast animals can switch between them. In tasks that involve simple reversal of motor response or sensorimotor mappings, monkeys usually use a small number of trials to switch between rules [81, 82, 83]. On the other hand, for WCST with more than two rules, as is usually used for humans, subjects typically use more trials to switch to the new rule [1, 84].

There are other reasons that may contribute to the suboptimality of behavior during rule switching, including random exploration [84], poor sensitivity to negative feedback [84], integration of reward history across multiple trials [85, 80, 12, 86], the gradual update of the value of the counterfactual rule [87] or the cost of cognitive control [88]. Neuronal mechanisms on longer timescales such as synaptic mechanisms [89] may be required to produce the slow switching behavior.

### Other limitations of the work

The SST neurons in the PFC module of our networks are targeted by the local VIP neurons. However, since the VIP neurons in our setup do not receive any excitatory input, their inhibition onto SST neurons is always released. Indeed, inhibiting the PFC VIP neurons in our the networks did not significantly affect task performance (Supplementary Figure 5f). In reality, VIP neurons in the PFC module receive long-range connections from the mediodorsal thalamus [90] which carries information about task context [91]. In addition, their activity is also strongly modulated by negative reinforcement [15] as well as arousal level [92], which suggest that they receive long-range projections from subcortical regions such as ventral tegmental area, locus coeruleus [93] and the raphe nucleus [94]. Including realistic models of these subcortical regions into the current model is an interesting direction for future work.

In conclusion, our approach of incorporating neurobiological knowledge into training RNNs can provide a fruitful way to build circuit models that are functional, high-dimensional, and reflect the heterogeneity of biological neural networks. In addition, dissecting these networks can make useful cross-level predictions that connect biological ingredients with circuit mechanisms and cognitive functions.

## Acknowledgements

This work was supported by James Simons Foundation Grant 543057SPI, the National Institutes of Health grant R01MH062349, and the ONR grant N00014-23-1-2040. YL thanks (alphabetically) Aldo Battista, Mark Buckley, Sage Chen, Shuo Chen, Vishwa Goudar, Kenneth Kay, Haohong Li’s lab, Jianguang Ni’s lab, Yu Qi’s lab, Yi Sun’s lab, Lucas Tian, Bo Shen, Xiaohan Zhang and all members of Xiao-Jing Wang’s lab for helpful discussions and comments on the manuscript.

## Methods

### Model setup

The RNN consists of two bidirectionally-connected modules, the PFC module and the sensorimotor module. Each module consists of 70 excitatory neurons and 30 inhibitory neurons. Each excitatory neuron has 2 dendritic compartments. The inhibitory neurons are evenly divided into three types: PV, SST and VIP. Different types of neurons have different connectivity, inspired by experimental findings [78, 95]: PV neurons target the somatic compartment of excitatory neurons and other PV neurons, SST neurons target the dendritic compartment of excitatory neurons as well as PV and VIP neurons, and VIP neurons target SST neurons. Excitatory neurons target other excitatory neurons, PV and SST neurons. The connection strength between all other types of neurons were fixed at zero throughout training.

Only excitatory neurons send long-range projections to other modules. The long-range projections from the sensorimotor module to the PFC module target the dendritic compartment of the excitatory neurons and the PV neurons. This is inspired by the experimental evidence that PV neurons mediate feedforward inhibition [14]. The long-range top-down projections from the PFC to the sensorimotor module target the dendritic compartments of the excitatory neurons and all three types of inhibitory neurons. Finally, external inputs to both modules target the dendritic compartment of excitatory neurons and PV neurons.

The dynamics of the somata of the excitatory neurons in the RNN are described by

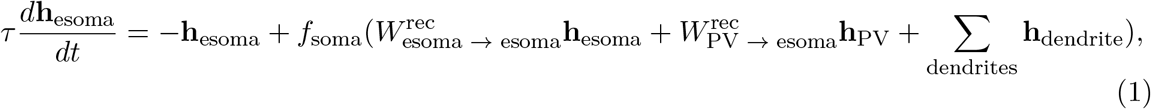

where *τ* = 100 ms, *dt* = 10 ms. Somata of excitatory neurons in both the sensorimotor and PFC modules obey the same equation. Here “esoma” stands for the soma of excitatory neurons. 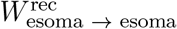 and 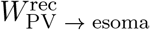 represent the connectivity matrix between the soma of excitatory neurons and from the local PV neurons to the soma of excitatory neurons, respectively. *h*_esoma_ and *h*_*P V*_ are the activity of the soma of excitatory neurons and PV neurons. *h*_dendrite_ is the activity of the dendritic compartment. *f*_soma_ is the somatic nonlinear activation function which was modeled as a rectified linear function:

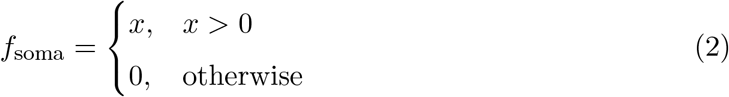

The dendritic activity is a nonlinear function of the excitatory and inhibitory inputs.

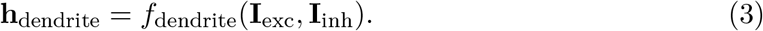

**I**_exc_ is the total excitatory input to the dendrite. It consists of long-range inputs from the input neurons (neurons that encode the feedback for the PFC module and neurons that encode the stimulus for the sensorimotor module) as well as the long-range input from the excitatory neurons in the other module. **I**_exc_ = **I**_in_ + **I**_cross-module_. **I**_inh_ is the inhibitory input to the dendrite from the local SST neurons. **I**_inh_ = **I**_SST→edend_. Here “edend” stands for the dendrite of excitatory neurons. The functional form of *f*_dendrite_ is described in the next section.

The inhibitory neurons are modeled as standard point neurons. Different types of inhibitory neurons receive different input connections. In the sensorimotor (SM) module, the dynamics of PV neurons are described by

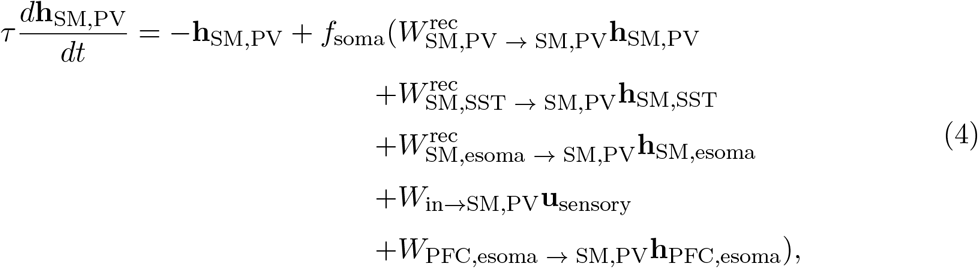

where 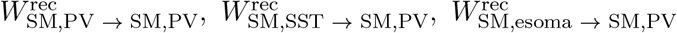 are the connection weight matrices between the PV neurons, from local SST neurons to the PV neurons, and from local excitatory neurons to the PV neurons, respectively. *W*_in→SM,PV_ is the input weight matrix to the PV neurons, and **u**_sensory_ is the input to the sensorimotor module that represents the features about the cards. *W*_PFC,esoma → SM,PV_*h*_PFC,esoma_ is the top-down connection weight matrix from the excitatory neurons in the PFC module to the PV neurons in the sensorimotor module.

For the SST neurons,

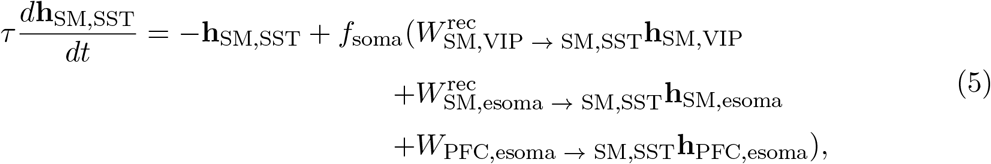

where 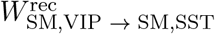 and 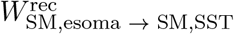are the connection weight matrices from local VIP neurons and excitatory neurons to the SST neurons, and *W*_PFC,esoma → SM,SST_*h*_PFC,esoma_ is the top-down connection weight matrix from the excitatory neurons in the PFC module to the SST neurons in the sensorimotor module.

For the VIP neurons,

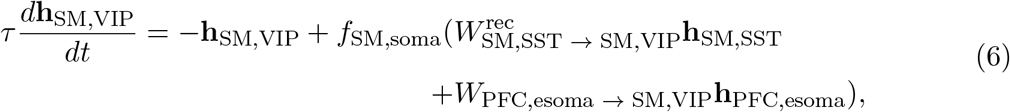

where 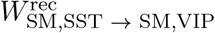 are the connection weight matrix from the local SST neurons to the VIP neurons, and *W*_PFC,esoma → SM,VIP_*h*_PFC,esoma_ is the top-down connection weight matrix from the excitatory neurons in the PFC module to the VIP neurons in the sensorimotor module.

The inhibitory neurons in the PFC module are described by similar equations, except only the PV neurons receive long-range bottom-up inputs from the sensorimotor module:

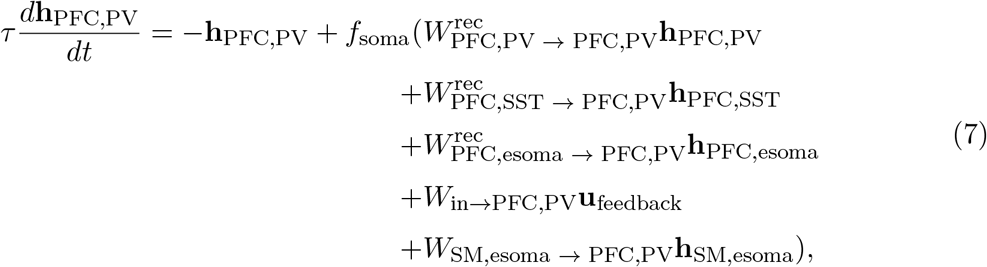

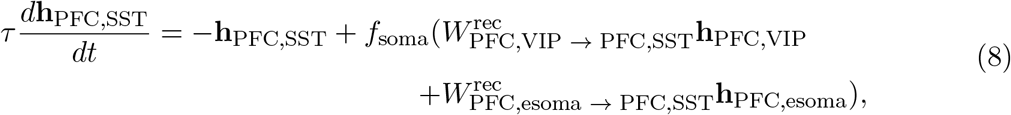

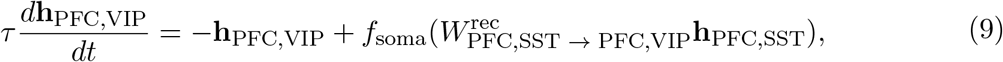

where **u**_feedback_ represents the external input to the PFC module about the feedback of the previous trial.

In practice, we used a mask matrix to enforce the connectivity between different cell types.

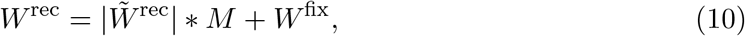

where 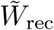 is the unconstrained connectivity matrix updated by the learning algorithm, *M* is a matrix consisting of 1, 0 and −1 depending on whether the corresponding connection is excitatory, inhibitory or nonexistent. *W*^fix^ implements the fixed coupling between the dendrite and the soma. * represents element-wise product.

Only the somata of excitatory neurons send output connections. The output in each module is generated via a simple linear readout:

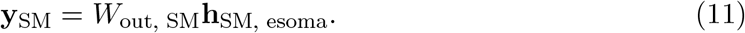

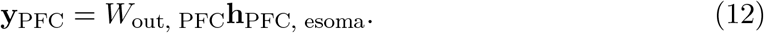

Both the input and output connection weights were constrained to be positive.

### Variations in the model hyperparameters

#### Dendritic nonlinearities

We trained models with two types of dendritic nonlinearities *f*_dendrite_ - subtractive and divisive. They are inspired by in-vitro and computational studies showing different types of inhibitory modulation on the dendritic activity depending on the location of inhibition relative to excitation [25]. Both types of dendritic nonlinearities are sigmoidal functions of the excitatory input. Under subtractive nonlinearity, as the inhibitory input increases, the turning point of the sigmoid function moves to larger values, consistent with the experimental observation when the inhibitory current is injected at the same location or more distal than the excitation [25]. For the divisive nonlinearity, the turning point of the sigmoid is not affected by the level of inhibition, but the saturating level of the sigmoid function decreases with the level of inhibition, consistent with the experimental observation when the inhibitory current is injected close to the soma [25].

The equations of the different dendritic nonlinearities are given by:

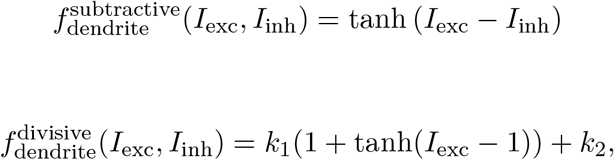

where 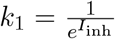 and *k*_2_ = −1 − tanh(−1). The form of the divisive dendritic nonlinearity was specified such that it is divisively modulated by *I*_inh_ (even when *I*_exc_=0), and that it is 0 only when both *I*_exc_ and *I*_inh_ are 0.

#### Initializations

The connectivity matrices were initialized either using a normal distribution with mean 0 and standard deviation 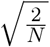 (where *N* is the total number of and recurrent units) or a uniform distribution between 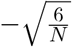 and 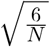.

### Sparsity of the SST→dendrite connectivity in the sensorimotor module

To study how the degree of dendritic branch-specific rule encoding in the sensorimotor module is affected by the sparsity of the connections from SST neurons to the dendrite of excitatory neurons, we varied this sparsity by fixing a fraction of randomly chosen connections to be 0 throughout training. The sparsity levels used were 0, 0.2, 0.4, 0.6 and 0.8.

### Random seeds

For each combination of the hyperparameter configuration introduced above (except the sparsity), we trained models using 50 random seeds for Pytorch (other random seeds were fixed). For each sparsity level other than 0, we trained models using 10 random seeds for Pytorch.

#### Task

The networks were trained on an analog of the Wisconsin Card Sorting Test (WCST) used for monkeys [23, 10, 11]. Each trial starts with the presentation of a “reference card” for 500 ms, after which three “test cards” appear around the reference card for 500 ms. Each card contains an object with a specific color (blue or red) and shape (circle or triangle). Among the three test cards, one of them matches the color of the reference card, another one matches the shape of the reference card, and the third card matches neither feature of the reference card. Depending on the rule (color or shape), the location where the test card has the same color or shape feature as the reference card should be chosen. The choice should be made during the 500 ms when both the reference card and the test cards are presented. At the end of this period, a feedback signal is presented for 100 ms, indicating whether the choice is correct or incorrect. This is followed by a 1 second inter-trial interval. The task rule switches after a random number of trials, without informing the network.

Therefore, the network inevitably makes an error for the first trial after the rule switch since it has not yet received the information that the rule has switched. The network should then adjust its behavior to the new rule by utilizing the feedback signal.

### Representation of inputs and outputs

Each card is represented as a four-dimensional binary vector, where different entries represent the presence of the two colors and shapes. The feedback input is a two-dimensional one-hot vector, where the two entries represent positive and negative feedback. The target output for the sensorimotor module is a three-dimensional one-hot vector, where each entry represents one response location on the screen. This target is non-zero only during the 500 ms response period when both the reference card and the test cards are presented. The target output for the PFC module is a two-dimensional one-hot vector, where each entry represents one rule. This target is non-zero during the entire trial.

### Training method

During training, the networks ran continuously across 20 consecutive trials with 3 random rule switches. Importantly, the network dynamics were not reset during the intertrial interval. The loss function was aggregated across the 20 trials.

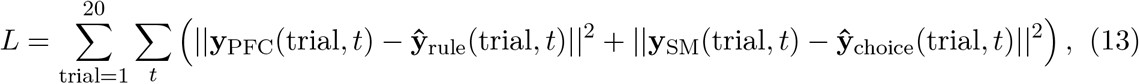

where **y**_PFC_(trial, *t*) and **y**_SM_(trial, *t*) are the activity of the readout neurons for the PFC and sensorimotor module at time *t* in a given trial, respectively. **ŷ**_rule_(trial, *t*) is the target output for the PFC module which represents the rule of the current trial. It is a binary vector of dimension 2 where each entry represents one rule. The activation of the entry that represents the correct rule is 1 throughout the entire trial. **ŷ**_choice_(trial, *t*) is the target output for the sensorimotor module which represents the correct choice for the current trial. It is a binary vector of dimension 3 where each entry represents one of the three response locations. The activation of the entry that represents the correct choice is 1 during the response period (500 ms when both the reference card and the tests card are shown).

The standard backpropogation through time algorithm [96] with the Adam optimizer [97] was used to update all connection weights.

We also used curriculum learning to speed up training. Initially, the stimulus, choice and outcome of the previous trial were all provided to the PFC module as input. This way all the information needed to perform the current trial was contained within the input, and the networks did not need to memorize past trials. During the training phase, the network performed 20 consecutive trials with 3 random rule switches, therefore the maximum performance was 85%. When the training performance reached above 65%, we started testing the network on longer trial sequences (200 consecutive trials with 10 rule switches). The maximum performance during testing was 95%. If the networks reached on average 90% performance during the recent 5 tests, the input about the previous stimulus was removed. When the networks reached 90% performance again, the information about the previous choice information was removed. The networks were then trained until they reached 90% performance.

Lower performance criteria was used for the model trained using early stopping (Figure 1f). In particular, curriculum training advanced to the next stage when the testing performance reached 80%.

### Single neuron selectivity metric

The selectivity index (SI) for rule is defined as

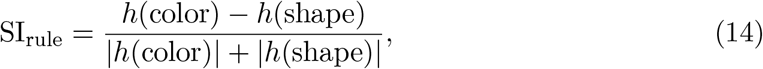

where *h*(color) and *h*(shape) represent the trial-averaged single neuron activity during color rule and shape rule, respectively. Neural activity was first averaged over the inter-trial interval before further being averaged across trials.

The error selectivity is defined similarly

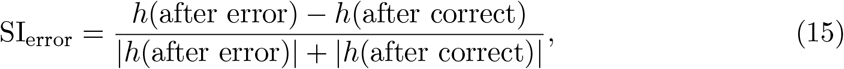

where *h*(after error) and *h*(after correct) are the mean single neuron activity after error and correct trials, respectively. Neural activity was first averaged across the feedback presentation and inter-trial interval periods before being averaged across trials.

The selectivity for response location is defined as

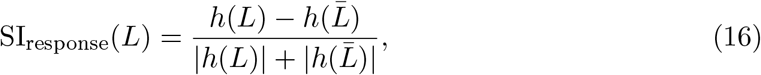

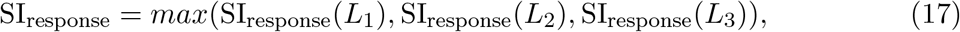

where *h*(*L*) represents the mean single neuron activity during trials where the network chooses response location *L*. 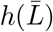 represents the mean activity across trials when the choice of the network is not location *L*. Therefore for each neuron we can compute three selectivity indices, one for each response location. The maximum of the three indices was taken as the response selectivity of that neuron and was used for Figure 5b. This selectivity index ranges from 0 to 1. We included neural activity during the response period when computing this selectivity index.

Neurons that prefer color/shape rule were further divided according to their preferred shared feature. The selectivity for the shared feature is defined as

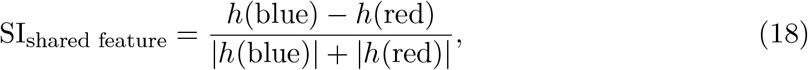

for neurons that prefer the color rule, and

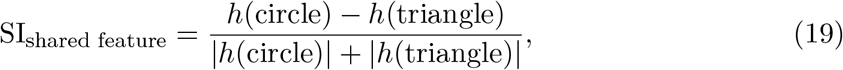

for neurons that prefer the shape rule. Here *h*(blue), *h*(red), *h*(circle), *h*(triangle) represent the mean activity of a neuron across trials when the reference card is blue, red (when the current rule is color), circle or triangle (when the current rule is shape). We included neural activity during the response period when computing this selectivity index.

### Classification criteria for different neuronal populations

Each neuron in the PFC module was classified as a “rule neuron” if the absolute value of its rule selectivity was greater than 0.5 and the absolute value of its error selectivity was smaller than 0.5. The rest of the neurons were classified as “error neurons” if their error selectivity was greater than 0.5. Error neurons with greater mean activity during the color rule trials that follow an error trial were defined as error x color rule neurons, and the other error neurons were defined as error x shape rule neurons.

Each neuron in the sensorimotor module was assigned with a preferred rule, response location and shared feature according to the condition during which it has the highest activity. There was no threshold for this classification. Neurons with zero activity during all trials were excluded from the analyses.

#### Connectivity bias

The connectivity bias (CB) was defined as the difference in the average connection weight between different sub-population of neurons. A positive value indicates an agreement with the simplified circuit diagram (Figures 3c, Figure 6h). For example, the connectivity bias from the PFC PV neurons to the PFC excitatory neurons is given by

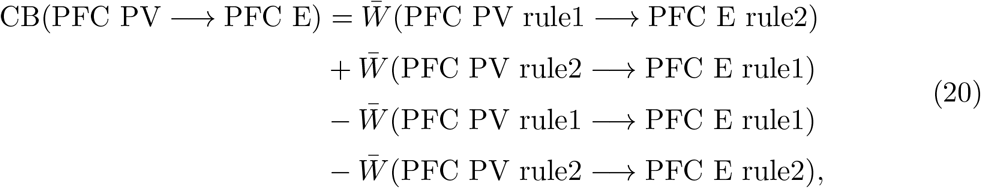

where for example 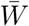 (PFC PV rule1 → PFC E rule2) represents the average (unsigned) connection strength from the PFC PV neurons that prefer rule 1 to PFC excitatory neurons that prefer rule 2. Here rule 1 refers to color rule and rule 2 refers to shape rule.

The other connectivity biases were defined analogously.

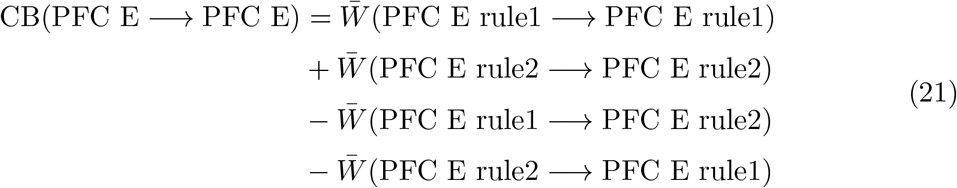

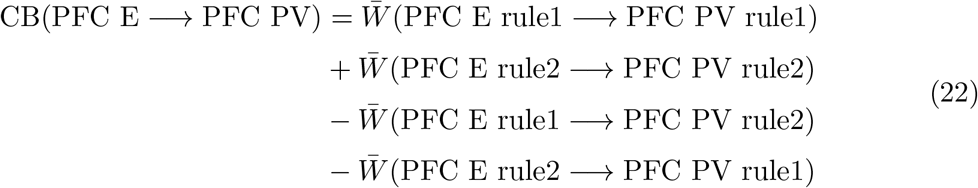

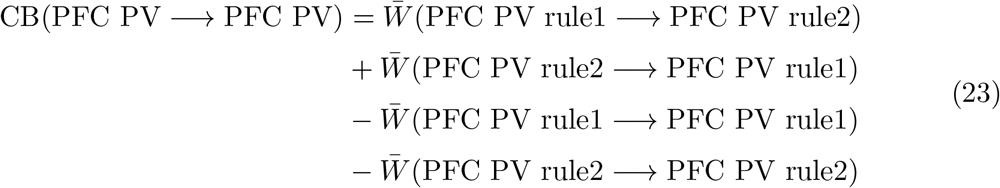

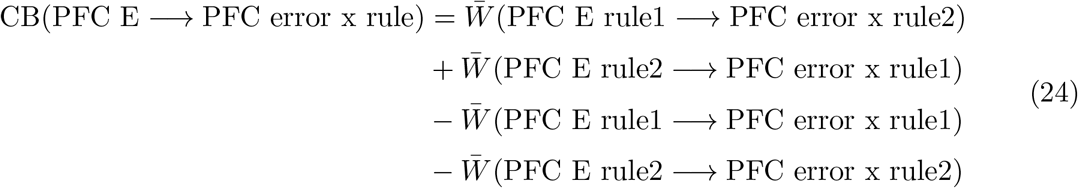

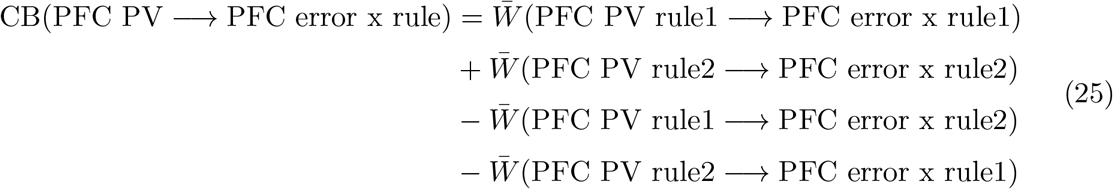

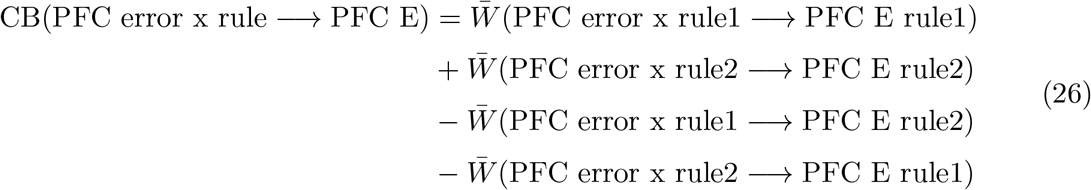

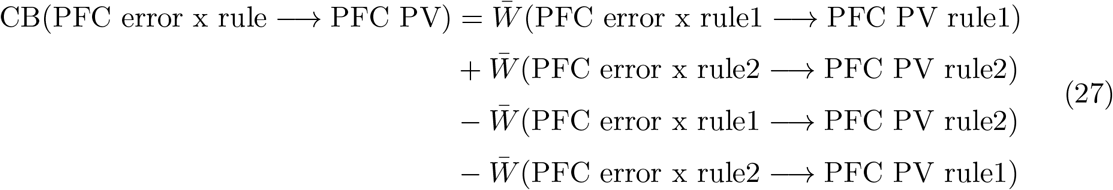

The connectivity biases between SST neurons and excitatory neurons in the PFC module (Supplementary Figure 5a-d) was defined similarly as those for PV neurons.

The connectivity biases between the different response location-selective populations in the sensorimotor module (SM) are defined as

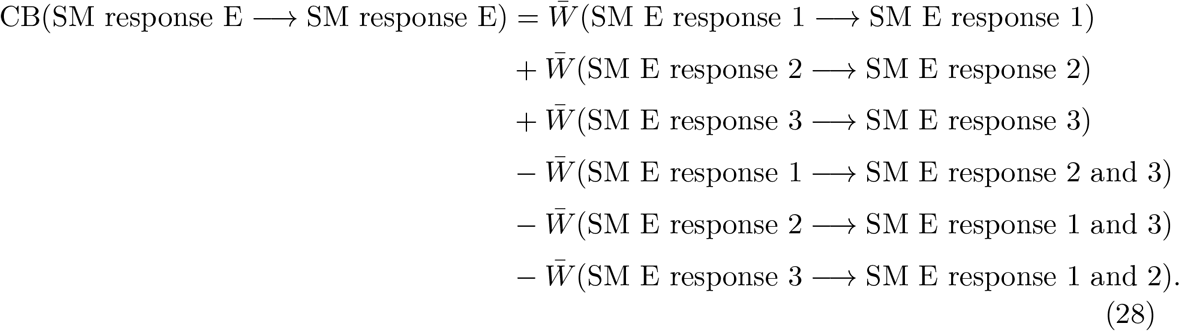

In the last equation, for example, 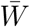 (SM response 1 → SM response 2 and 3) represents the mean connection strength from excitatory neurons in the sensorimotor module that prefer response location 1 to those that prefer response locations 2 and 3.

The other connectivity biases were defined similarly

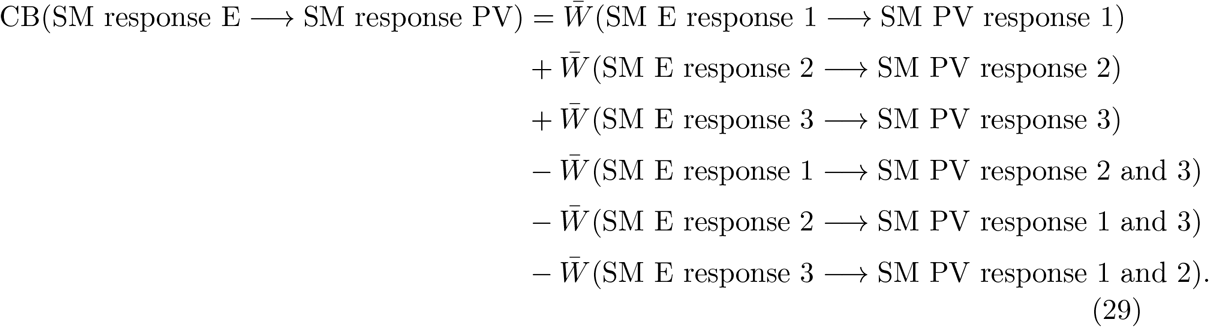

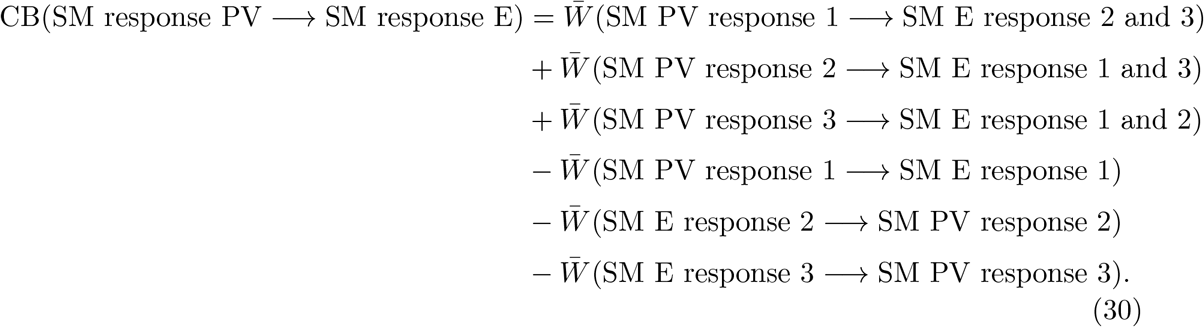

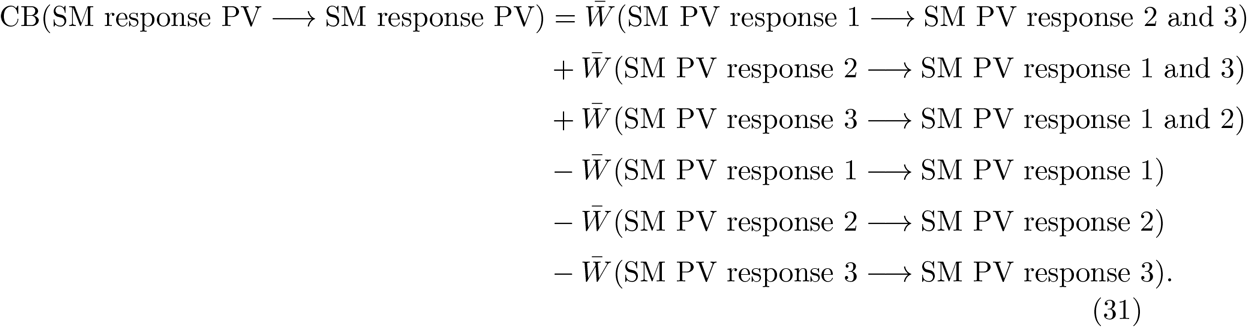

The connectivity biases between the different rule-selective populations in the sensorimotor module are defined as

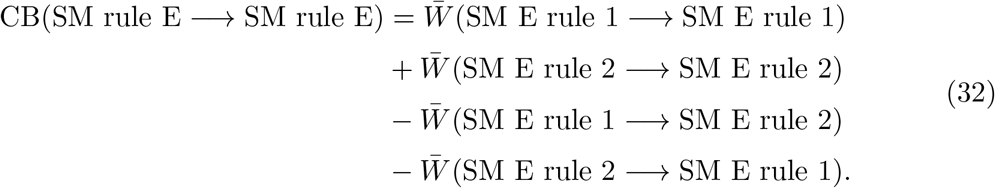

The other connectivity biases were defined similarly

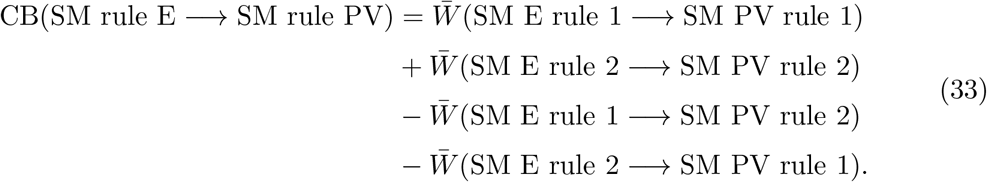

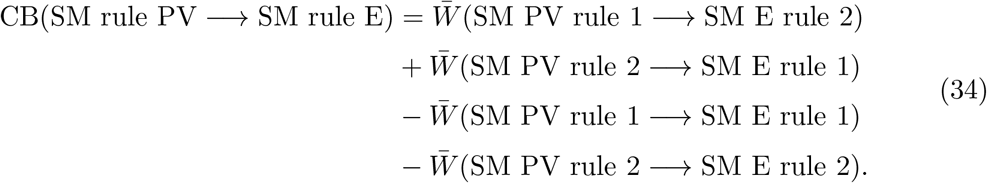

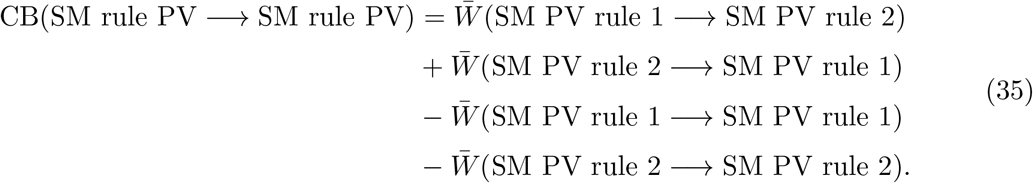

The connectivity biases between the different shared feature-selective populations in the sensorimotor module are defined similarly. For the populations selective for the two colors

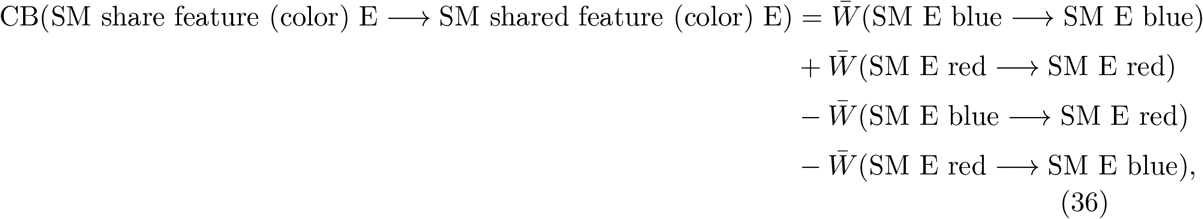

where for example 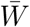 (SM E blue → SM E blue) is the average connection strength within the neural population selective for the shared feature blue.

The other connectivity biases were defined similarly

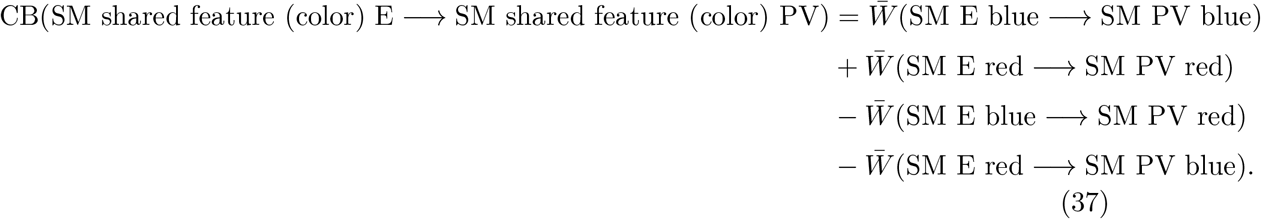

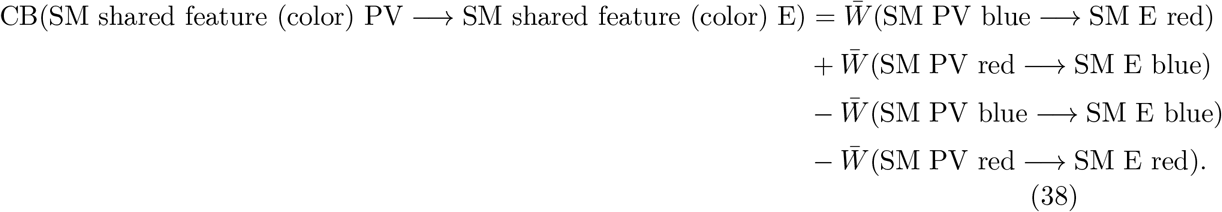

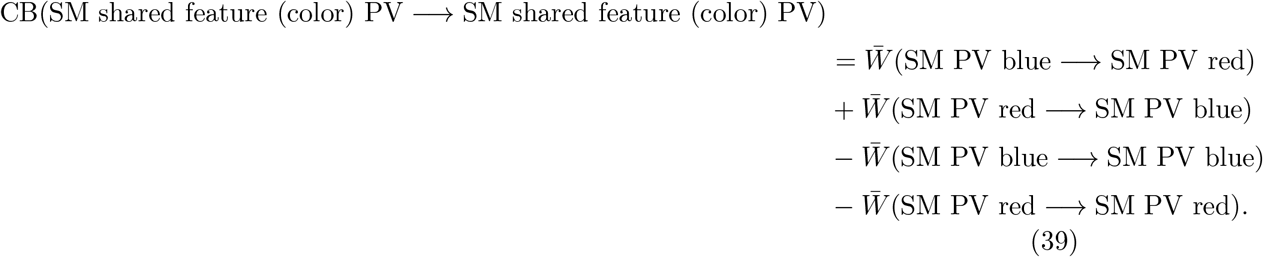

The connectivity biases between populations selective for different shared shapes were defined analogously by substituting blue and red with circle and triangle.

### Simulation of the optogenetic inhibition

Optogenetic inhibition was simulated by clamping the activity of neurons at 0 throughout the entire trial and the inter-trial interval.

### Principal angle between subspaces

The principal angle between two subspaces is a generalization of angle between lines and planes in Euclidean space to arbitrary dimensions [98]. It can be computed by iteratively finding pairs of unit length “principal vectors”, one from each subspace, that have the greatest inner product, subject to the condition that the principal vectors are orthogonal to all previous principal vectors [99].

In computing the principal angles between different rule-selective and response-selective subspaces, we first determined the dimensionality of the subspaces using the participation ratio [100]. Then the principal angles were computed using the “subspace_angles” function from the Python package Scipy. The largest principal angle was used.

To obtain a shuffled distribution, we first randomly and evenly split all trials belonging to a particular rule or response into two halves. Then, we generated two subspaces from neural trajectories during the two group of trials. A principal angle between these two subspaces was then computed for each rule/response. The angles were then averaged across all rules/responses to obtain a principal angle from shuffled data. This process was repeated 100 times to generate a distribution of principal angles from shuffled data.

### Assessing the strength of non-linear mixed selectivity

The extent to which neurons in the sensorimotor module encode the conjunction of stimulus and rule in a non-linear fashion was evaluated using the coefficient of determination of a linear regression model. To tease apart non-linear and linear mixed selectivity, we first fitted the mean activity of each neuron during response period using a set of regressors that represent either the rule or the stimulus alone:

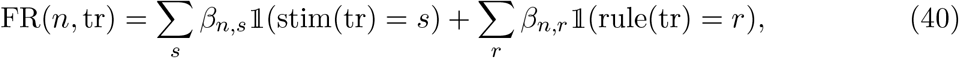

where FR(*n*, tr) is the mean firing rate of neuron *n* during the response epoch of trial tr. 𝟙 is the indicator function. For example, 𝟙 (stim(tr) = *s*) = 1 if the stimulus during trial tr is *s*, and it is 0 otherwise.

Then, another linear regression model was fitted on the residual activity unexplained by the linear regression model above, using the conjunction of rule and stimulus as regressors:

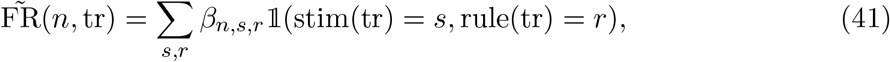

where 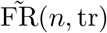 is the firing rate of neuron *n* during trial tr subtracted by the predicted firing rate from the model defined by Equation 40. The *R*^2^ value of this regression model was used to represent the strength of non-linear mixed selectivity.

**Acknowledgements**

This work was supported by James Simons Foundation Grant 543057SPI (X.-J.W.), the National Institutes of Health grant R01MH062349 (X.-J.W.), and the ONR grant N00014-23-1-2040 (X.-J.W.). We thank (alphabetically) Aldo Battista, Mark Buckley, Sage Chen, Shuo Chen, Vishwa Goudar, Kenneth Kay, Haohong Li’s lab, Jianguang Ni’s lab, Yu Qi’s lab, Yi Sun’s lab, Lucas Tian, Bo Shen, Xiaohan Zhang and all members of Xiao-Jing Wang’s lab for helpful discussions and comments on the manuscript.

## Data availability statement

All trained networks and pre-processed data used in this study in this study have been deposited at https://drive.google.com/drive/folders/17LqvmcBynX0a4OtUgnkODGJKK_JFRB2Q?usp=sharing

## Code availability statement

All computer code used to generate the results in this manuscript has been uploaded to https://github.com/liuyuue/BioRNN_WCST, archived in https://doi.org/10.5281/zenodo.10183167[101].

## Author Contributions Statement

Y.L. and X.-J.W. designed the research, discussed regularly throughout the project and wrote the manuscript. Y.L. performed the research.

---

**Supplementary Figure 1:**
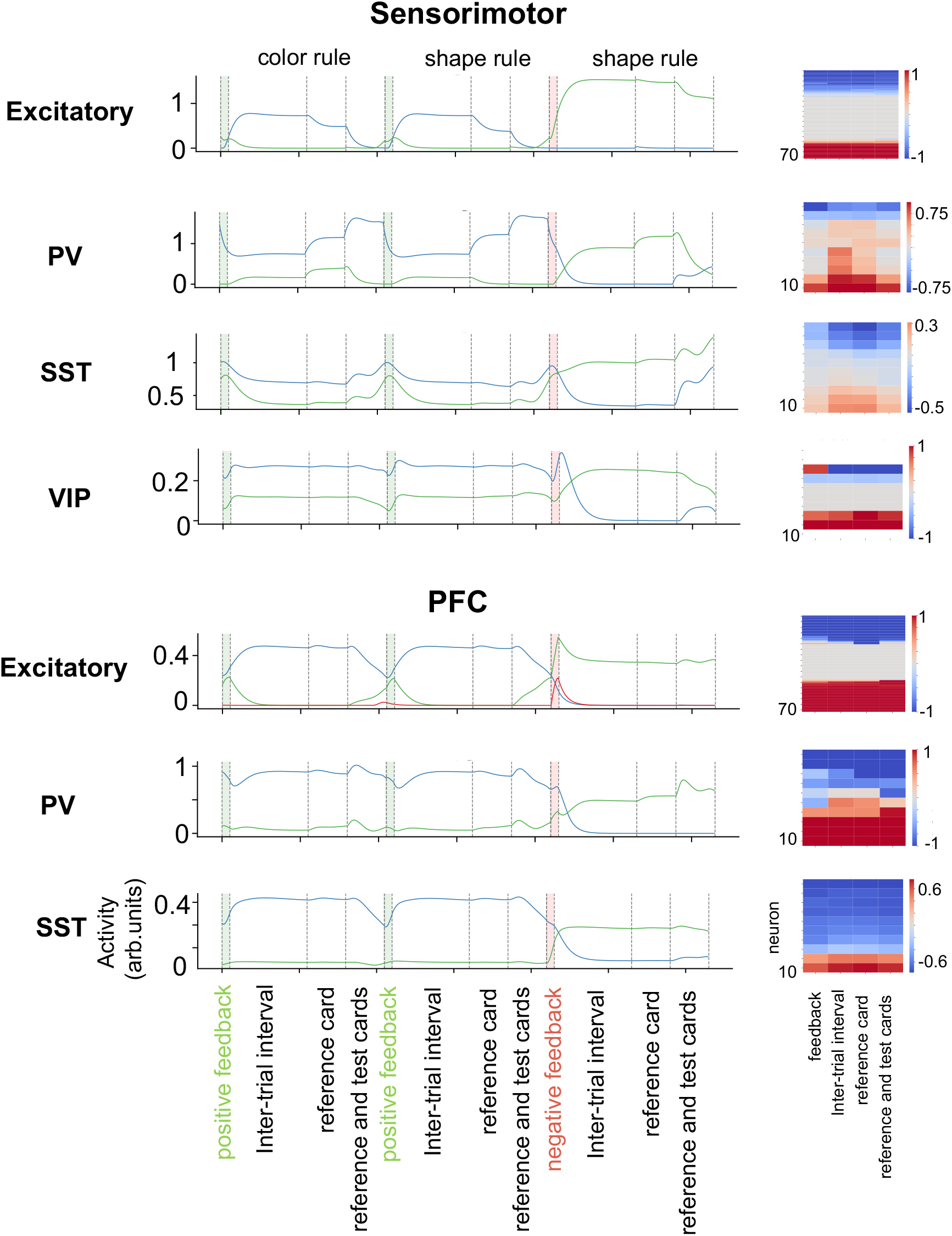
Activity of single neurons. **Left**: activity from example neurons across three consecutive trials with a rule switch. Blue and green traces represent neurons with higher activity during the color rule and shape rule respectively. Red trace for the PFC excitatory neurons represent neurons that are preferentially activated by negative feedback. **Right**: Rule selectivity across neurons and task epochs. VIP neurons in the PFC module do not receive any excitatory inputs, therefore are not shown.

**Supplementary Figure 2:**
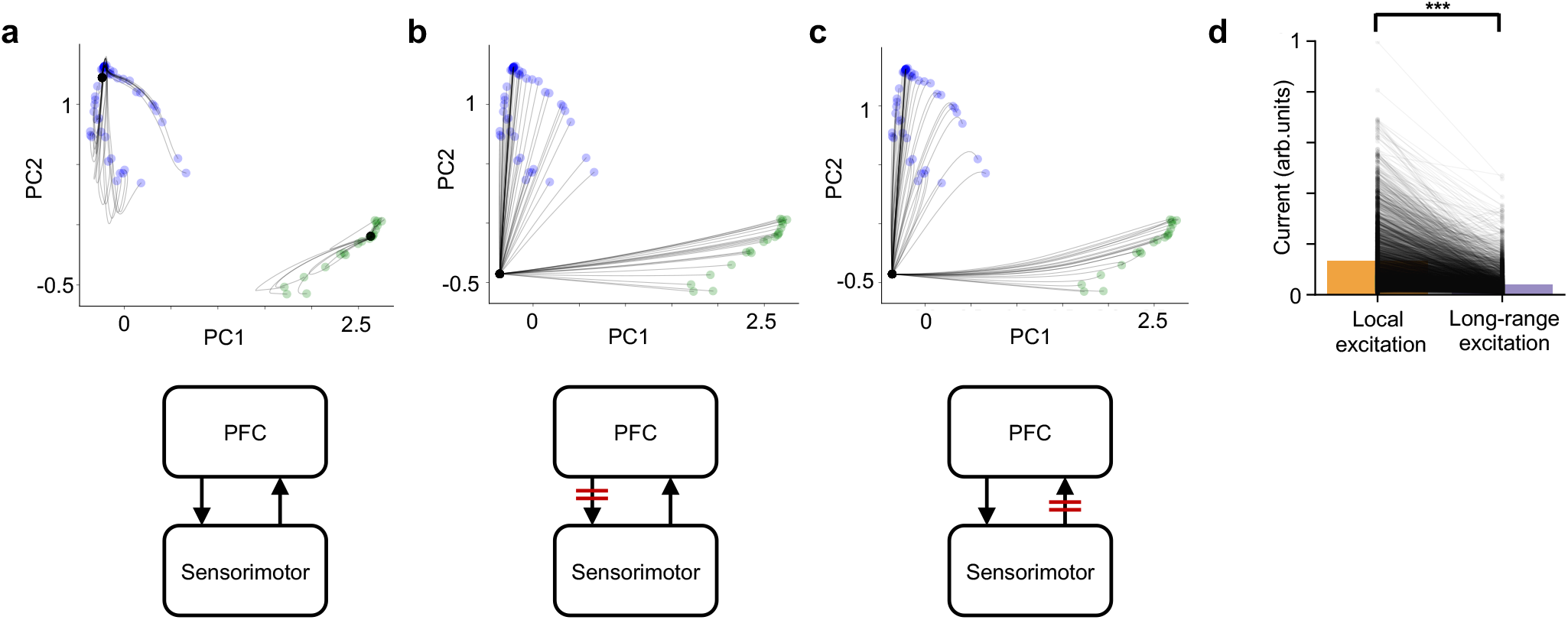
Two attractor states in the PFC module supported by inter-modular connections. The neural trajectories that represent the autonomous dynamics of the trained RNN with intact inter-modular connections (**a**), when the connections from the PFC module to the sensorimotor module were lesioned (**b**), and when the connections from the sensorimotor module to the PFC module were lesioned (**c**). The networks started from random time point during color rule (blue) and shape rule (green) trials. Black points represent the final states of the networks after 500 timesteps (5 seconds). **d**. Excitatory and PV neurons in the PFC module receive stronger local excitation than long-range excitation. Each line represents a neuron. Bars represent average across neurons. Result aggregated across all trained networks. One-sided Student’s t-test, *p* = 0.0, *n* = 4160 neurons.

**Supplementary Figure 3:**
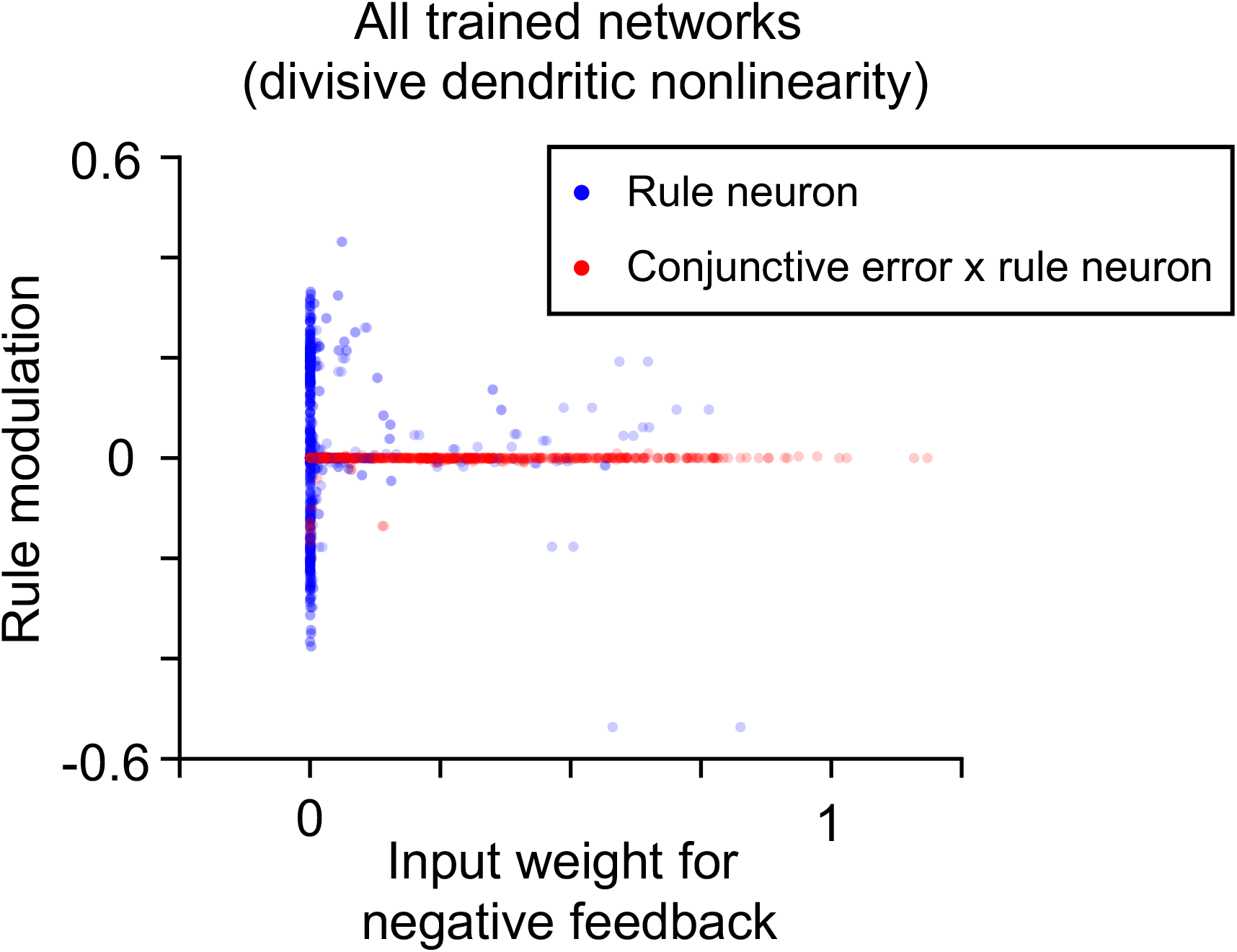
The emergence of two populations of excitatory neurons in the PFC module of networks with divisive dendritic nonlinearity. The rule modulation against input weight for negative feedback for all the rule neurons and conjunctive error x rule neurons in the PFC module of networks with divisive dendritic nonlinearity (c.f. Figure 2c).

**Supplementary Figure 4:**
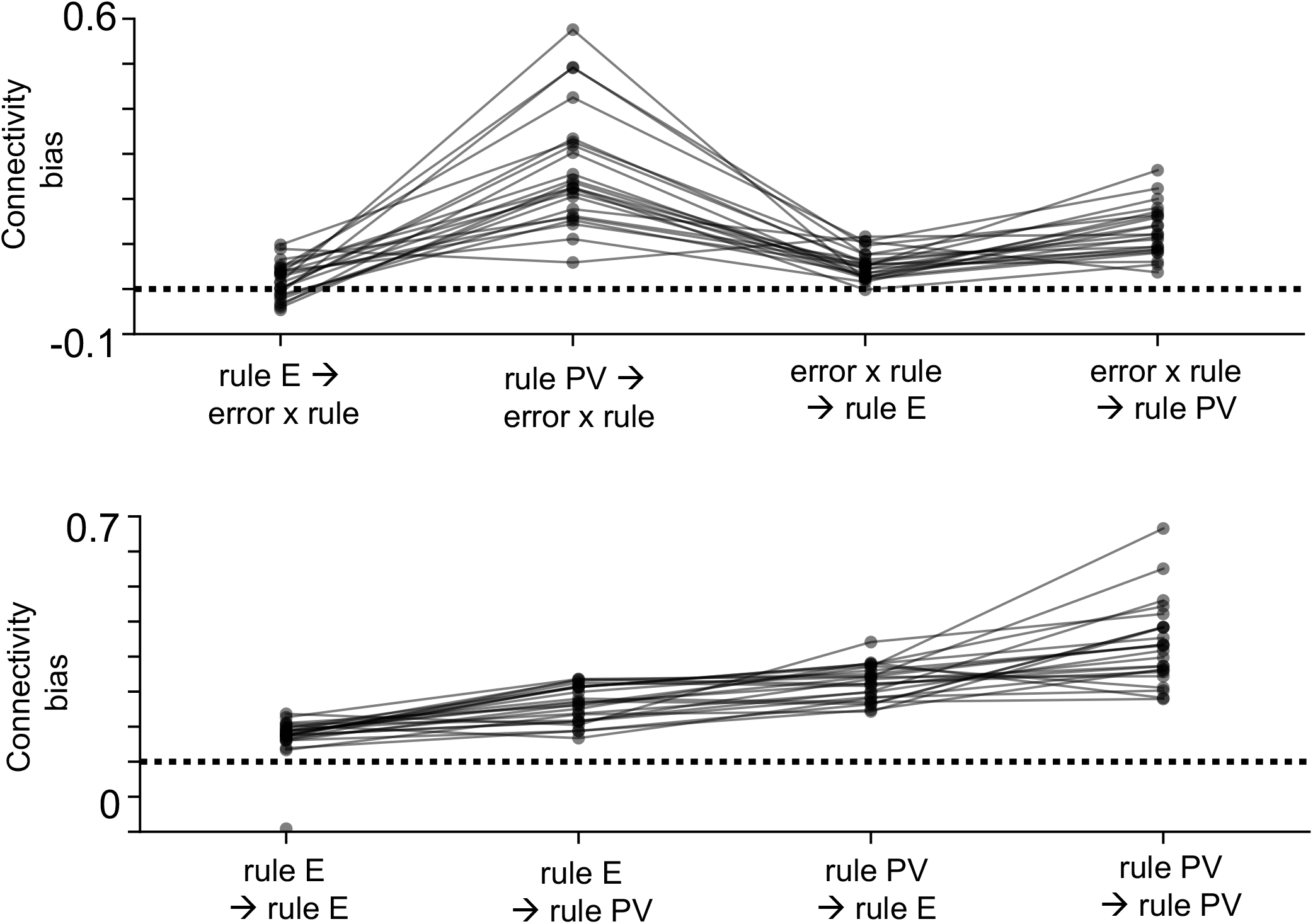
The connectivity biases between different subpopulations of excitatory and PV neurons in the PFC module of networks with divisive dendritic nonlinearity (c.f. Figure 3d).

**Supplementary Figure 5:**
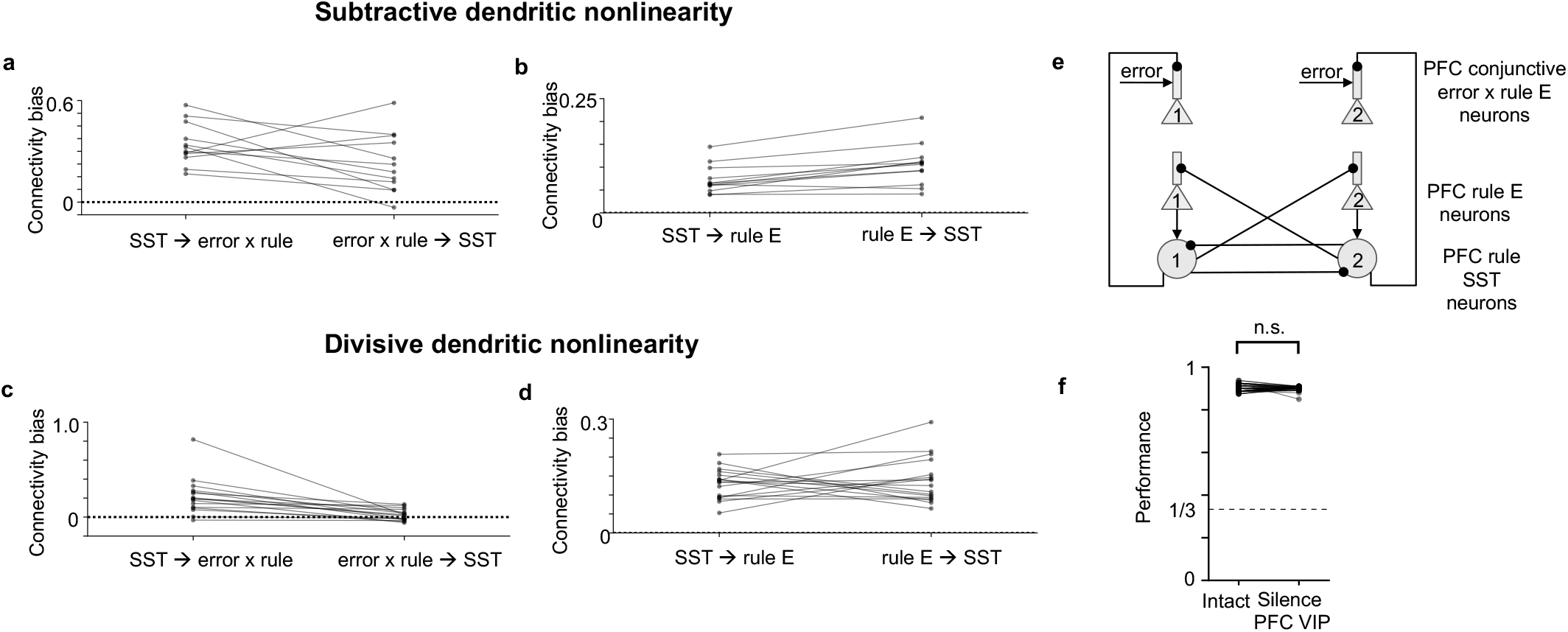
The connectivity structure between the excitatory neurons and SST neurons in the PFC module. a. The connectivity bias between the SST neurons and the conjunctive error x rule neurons in the PFC module, for networks with subtractive dendritic nonlinearity. **b**. The connectivity bias between the SST neurons and the rule neurons in the PFC module, for networks with subtractive dendritic nonlinearity. **c** - **d**. The same connectivity biases but for networks with divisive dendritic nonlinearity. **e**. The connectivity structure between the SST and excitatory neurons in the PFC module as revealed by **a**-**d**. Only the stronger connections are plotted. A similar connectivity pattern exists between SST and excitatory neurons as compared to the connectivity pattern between PV and excitatory neurons (c.f. Figure 3c). **f**. The performance is not affected by silencing VIP neurons in the PFC module (one-sided Student’s t-test, *p* = 0.49, *n* = 52 networks).

**Supplementary Figure 6:**
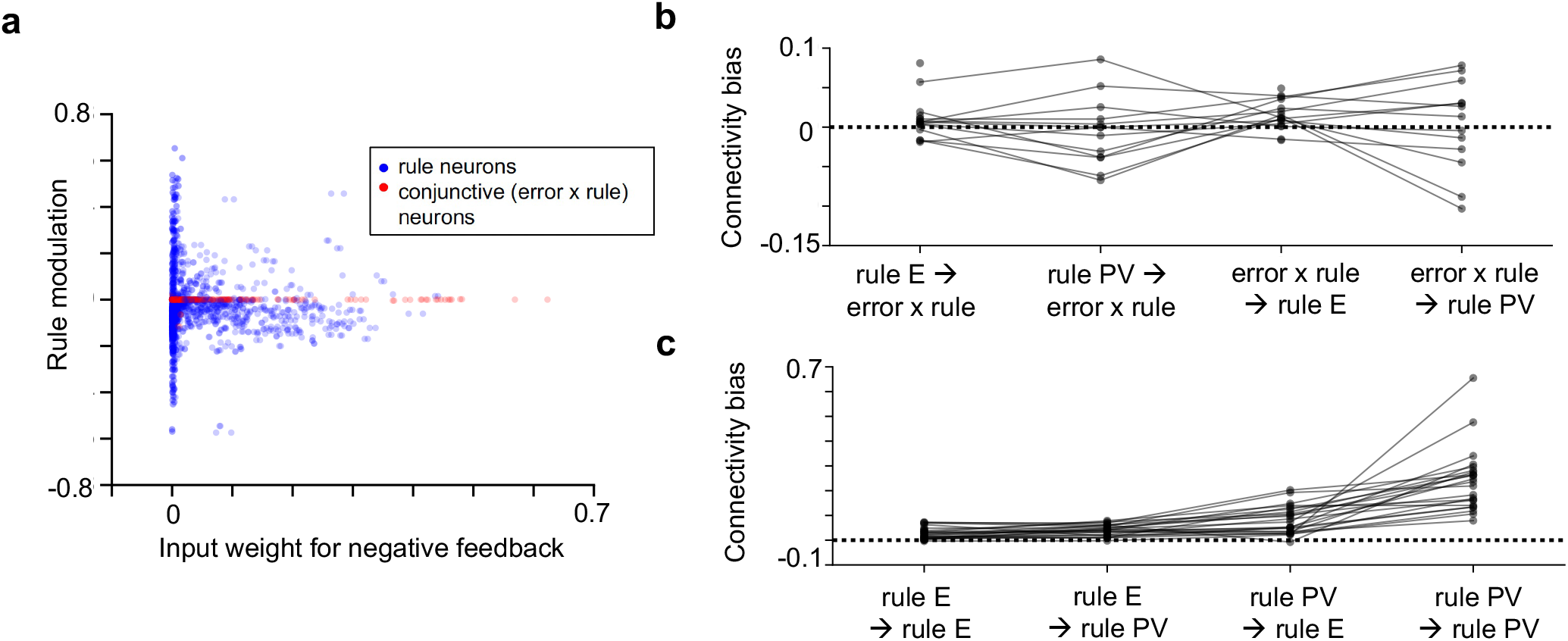
The connectivity biases for networks that shift between rules more gradually (c.f. Figure 1f). **a**. The rule modulation against input weight for negative feedback for all the rule neurons and conjunctive error x rule neurons in the PFC module of networks that switch rules more slowly. The separation of the two populations is weaker (compare with Figure 2c and Supplementary Figure 3) **b**. The connectivity biases between the error x rule neurons and rule neurons. **c**. The connectivity biases between rule neurons. The connectivity biases between error x rule neurons and rule neurons are not significantly greater than 0, whereas the ones between rule neurons are. This indicates that rule neurons form a winner-take-all attractor network, similar to the fast switching models (c.f. the lower panels of Figure 3d, Supplementary Figure 4), whereas the connections between the rule neurons and conjunctive error x rule neurons are less structured. The connectivity biases were only computed for models where the corresponding neuronal populations can be defined.

**Supplementary Figure 7:**
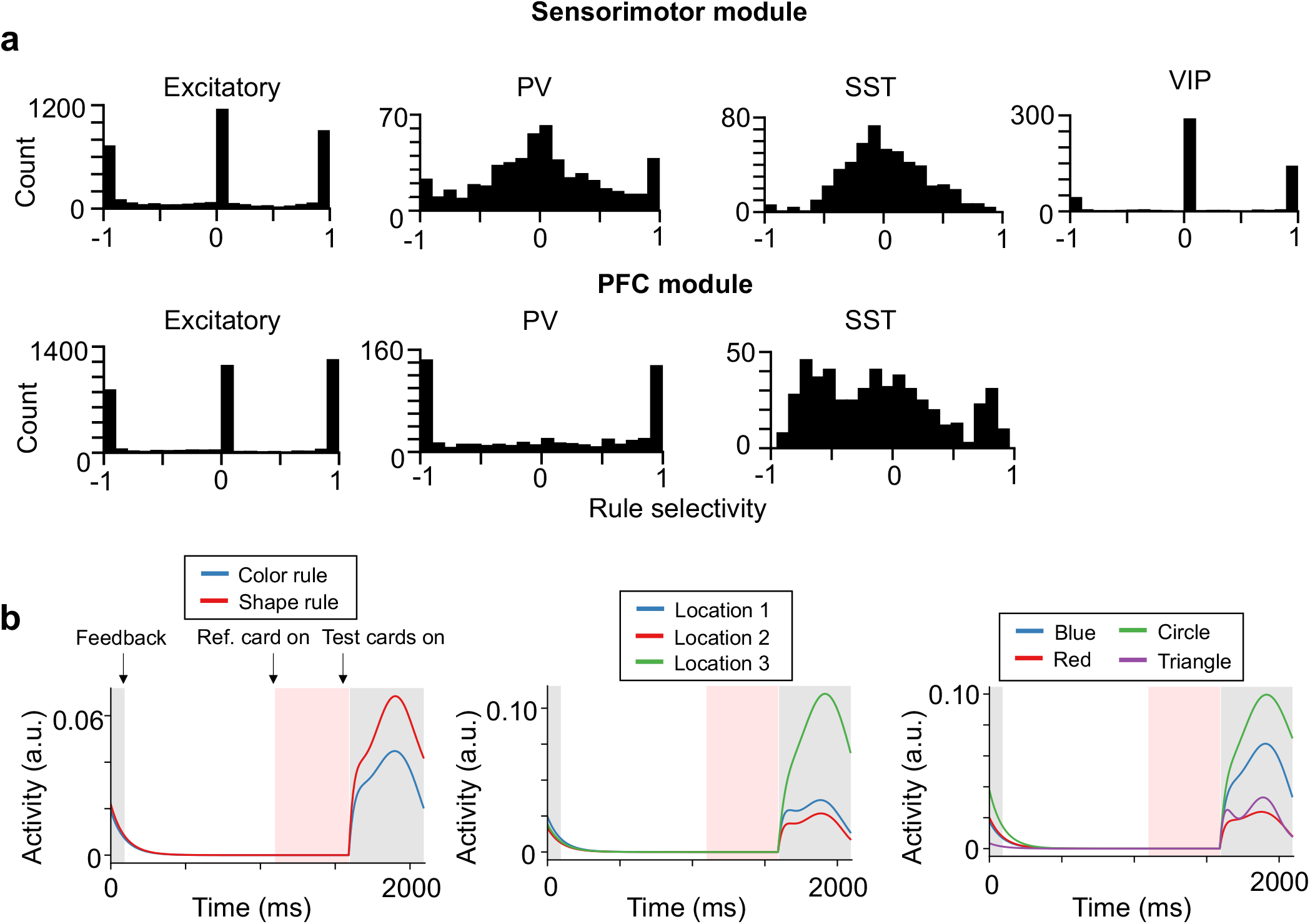
Single neuron feature selectivity in the PFC and sensorimotor modules. **a**. The distribution of rule selectivity across different cell types. Result is aggregated across all trained networks. Result for VIP neurons in the PFC module is not shown since they do not receive excitation. **b**. The trial-averaged activity of an example excitatory neuron in the sensorimotor module with preferred rule *R* = *color rule*, response location *L* = 3 and shared feature *F* = *circle*. Trials were sorted according to rule (left), response (middle) and shared feature (right).

**Supplementary Figure 8:**
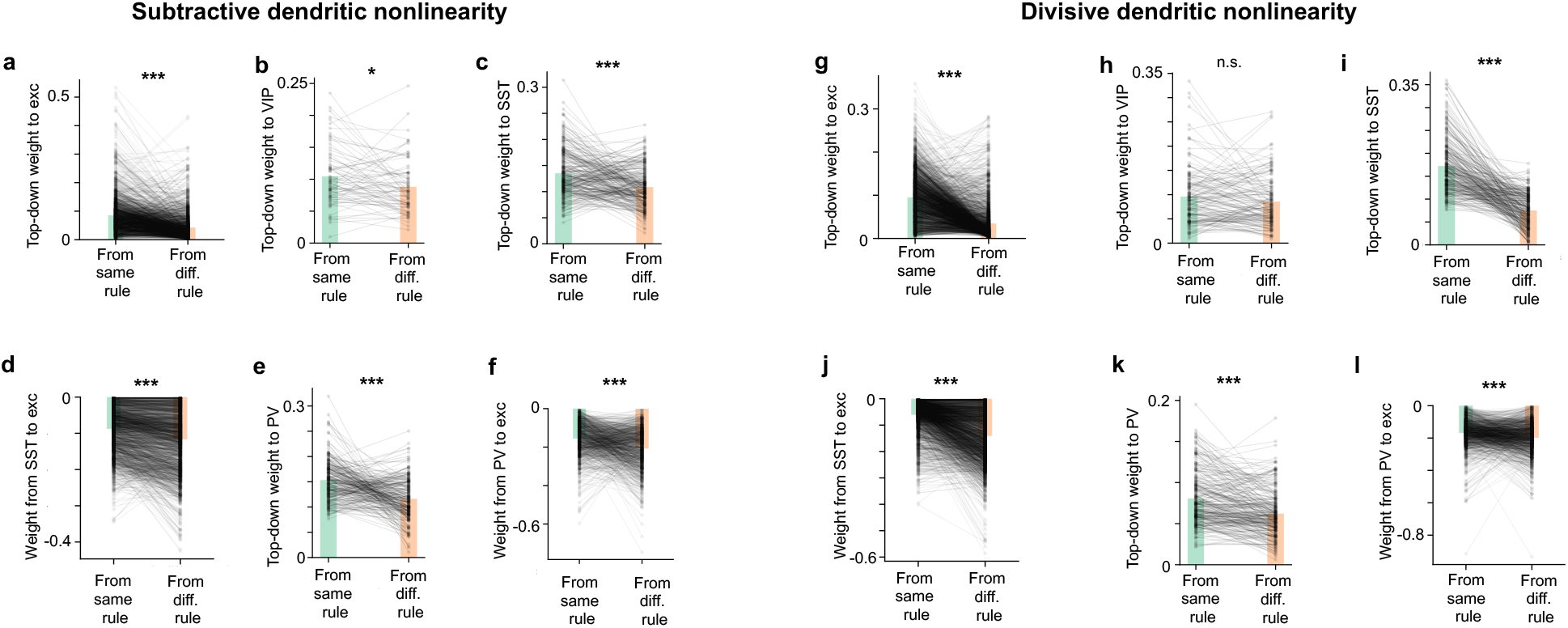
Structure in the top-down projections, across all networks. **a**. Each line represents the mean connection strength onto one dendritic branch of the excitatory neurons in the sensorimotor module, from the PFC excitatory neurons that prefer the same rule and the different rule. Bars represent the mean across neurons. PFC excitatory neurons project more strongly to the excitatory neurons in the sensorimotor module that prefer the same rule (c.f. Figure 4a). One-sided Student’s t-test, *p* = 1.6∗ 10^−124^, *n* = 2060 dendritic branches. **b**. The same data but for connections from excitatory neurons in the PFC module to VIP neurons in the sensorimotor module. One-sided Student’s t-test, *p* = 0.02, *n* = 79 VIP neurons (c.f. Figure 4b). **c**. The same data but for connections from excitatory neurons in the PFC module to SST neurons in the sensorimotor module. One-sided Student’s t-test, *p* = 2.3 ∗ 0^−12^, *n* = 250 SST neurons (c.f. Figure 4c). **d**. The same data but for connections from SST neurons to excitatory neurons in the sensorimotor module. One-sided Student’s t-test, *p* = 3.3∗10^−37^, *n* = 2060 dendritic branches (c.f. Figure 4d). **e**. The same data but for connections from excitatory neurons in the PFC module to PV neurons in the sensorimotor module. One-sided Student’s t-test, *p* = 3.8 ∗ 10^−23^, *n* = 250 PV neurons (c.f. Figure 4e). **f**. The same data but for connections from PV neurons to excitatory neurons in the sensorimotor module. One-sided Student’s t-test, *p* = 2.3 ∗ 10^−34^, *n* = 1030 excitatory neurons (c.f. Figure 4f). **g** - **l**. Same as **a** - **f** for networks with the divisive dendritic nonlinearity. **g**. One-sided Student’s t-test, *p* = 0.0, *n* = 2524 dendritic branches. **h**. One-sided Student’s t-test, *p* = 0.1, *n* = 148 VIP neurons. **i**. One-sided Student’s t-test, *p* = 3.0 ∗ 10^−79^, *n* = 260 SST neurons. **j**. One-sided Student’s t-test, *p* = 9.1 ∗ 10^−251^, *n* = 2060 dendritic branches. **k**. One-sided Student’s t-test, *p* = 9.6 ∗ 10^−12^, *n* = 260 PV neurons. **l**. One-sided Student’s t-test, *p* = 1.4 ∗ 10^−16^, *n* = 1262 excitatory neurons.

**Supplementary Figure 9:**
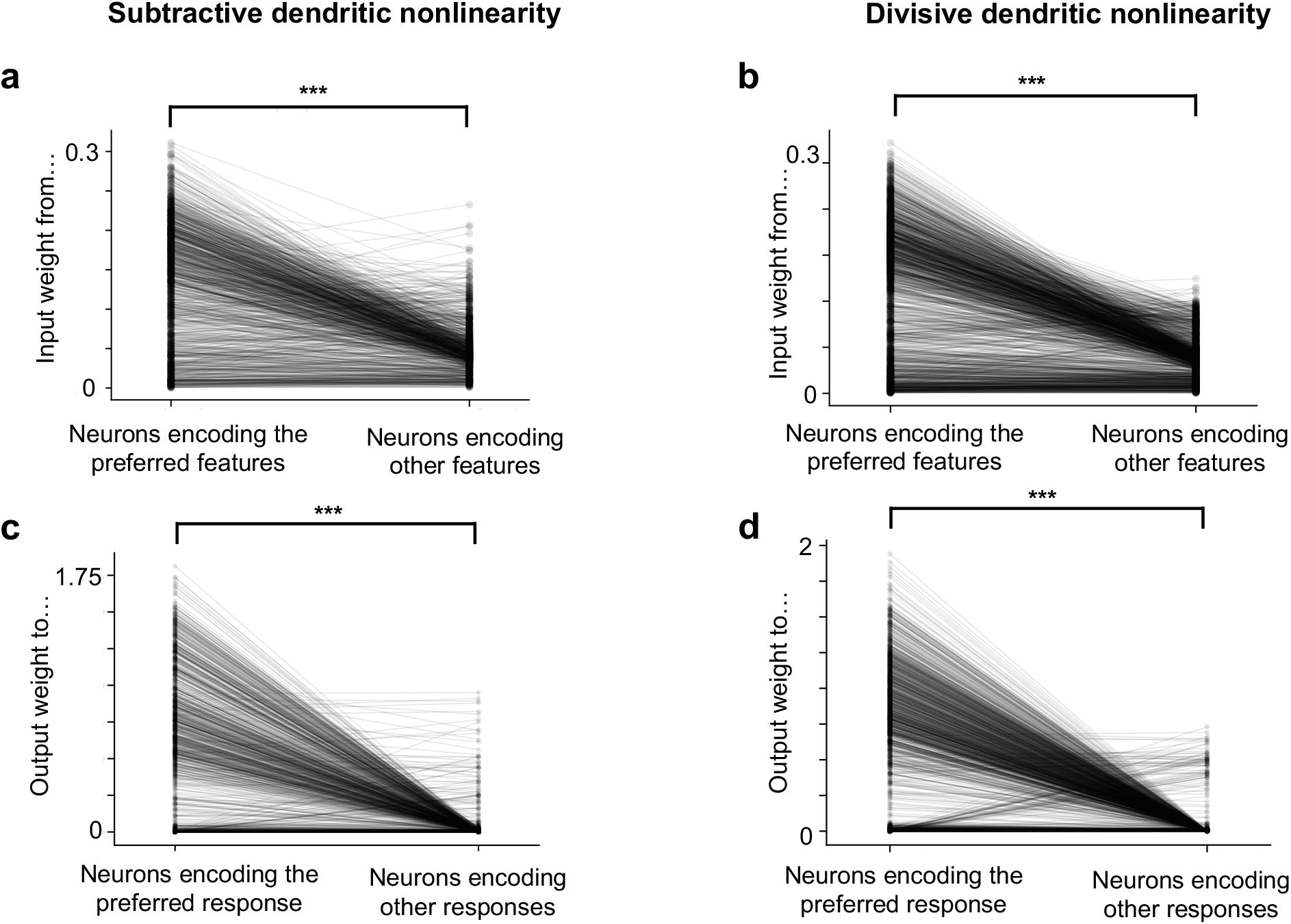
The structure in the input and output weights of the sensorimotor module. Data is aggregated across all trained networks with subtractive (**a, c**) and divisive (**b, d**) dendritic nonlinearity (c.f. Figure 5b, c). One-sided Student’s t-test was used for all panels. **a**: *p* = 4.4 ∗ 10^−189^, *n* = 1025 neurons. **b**: *p* = 1.1 ∗ 10^−270^, *n* = 1468 neurons. **c**: *p* = 4.5 ∗ 10^−218^, *n* = 1030 neurons. **d**: *p* = 6.2 ∗ 10^−308^, *n* = 1508 neurons.

**Supplementary Figure 10:**
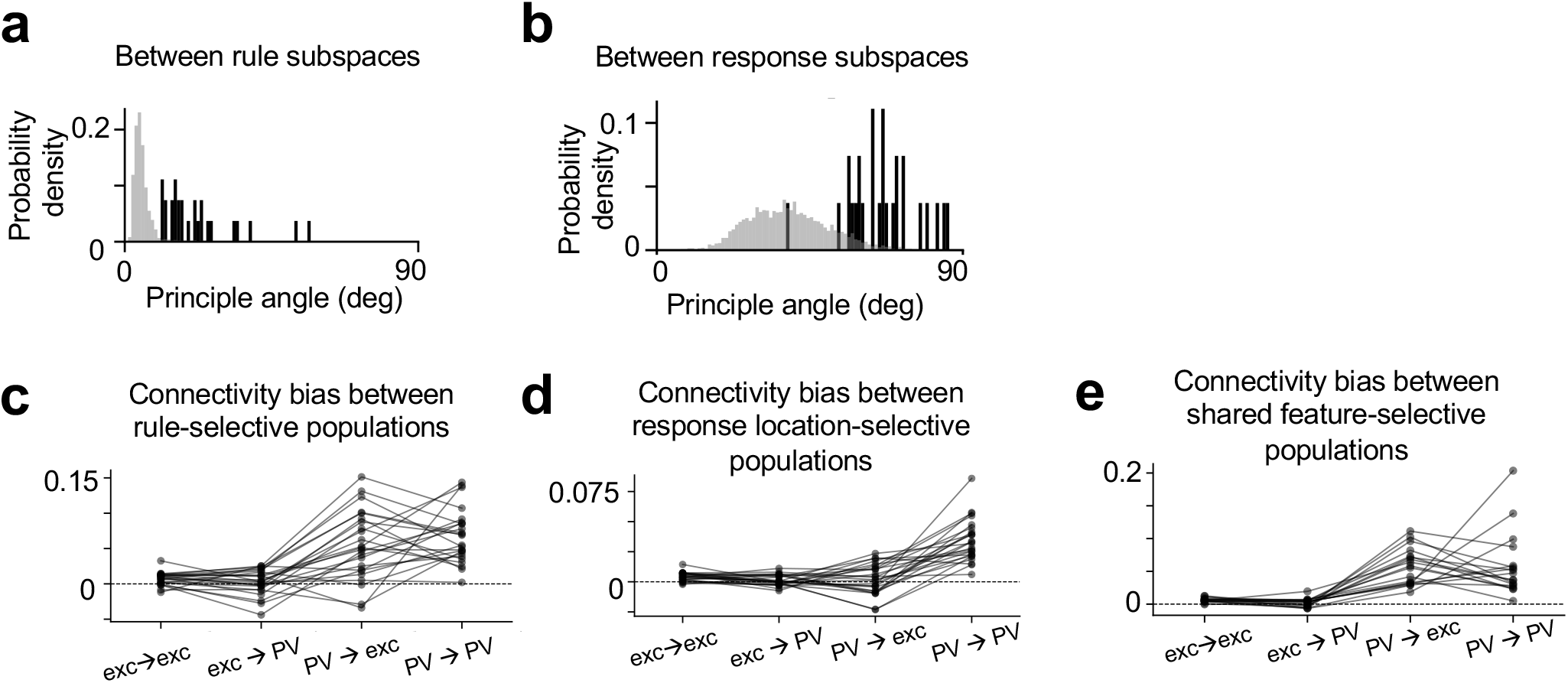
**a**-**b**. The principal angle between different rule and response subspaces for networks trained with divisive dendritic nonlinearity (c.f. Figure 6c-d). **c**-**e**. The connectivity bias between different subpopulations of excitatory and PV neurons in the sensorimotor module of networks with divisive dendritic nonlinearity (c.f. Figure 6e-g).

**Supplementary Figure 11:**
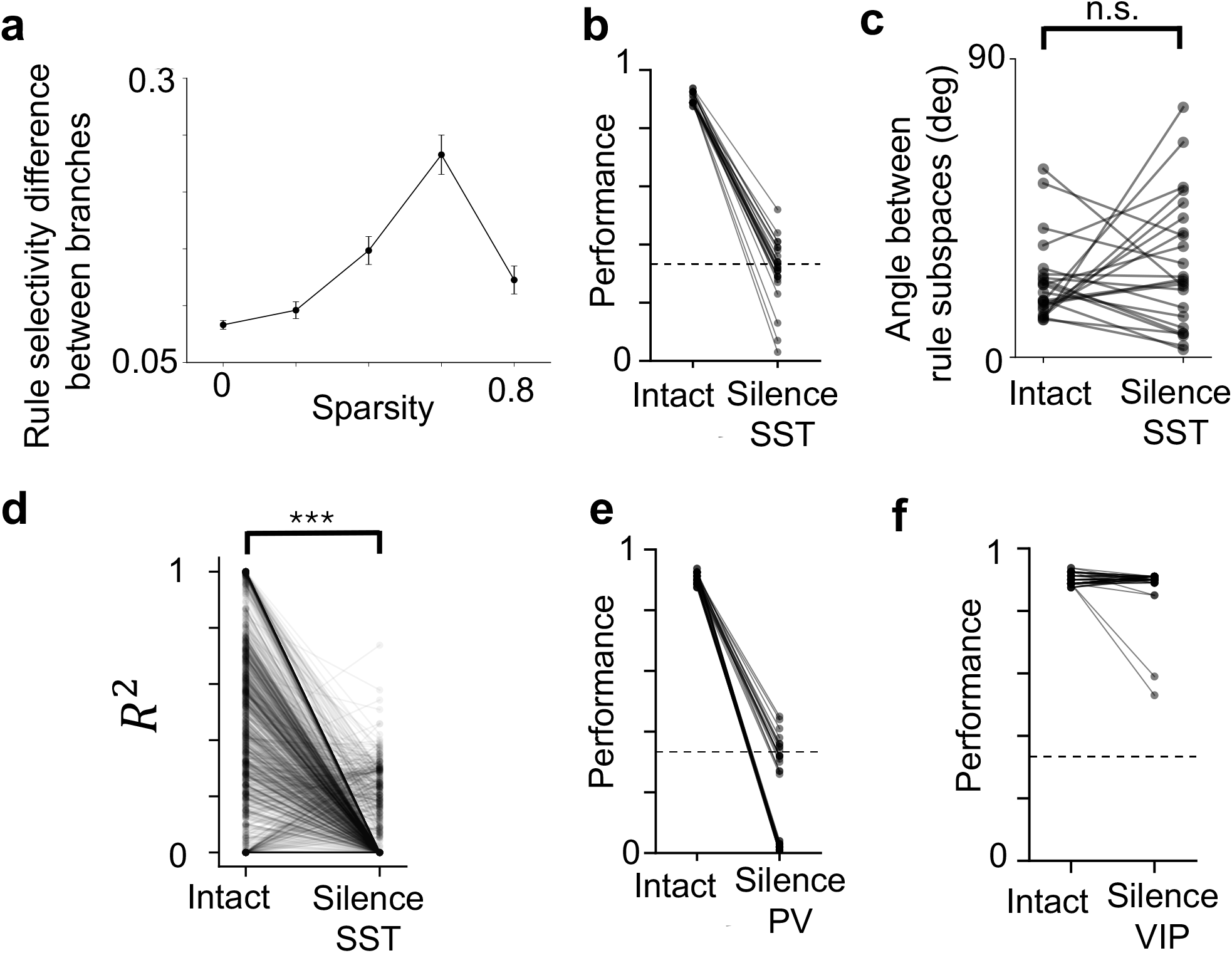
**a**. The relationship between the sparsity of the connection from SST neurons to the dendrites of excitatory neurons in the sensorimotor module and the degree of dendritic branch specific coding of the task rule, for networks with divisive dendritic nonlinearity (c.f. Figure 7d). Error bars represent standard error of the mean across *n* = 1960, 490, 350, 280, 350 pairs of dendritic branches for sparsity levels 0, 0.2, 0.4, 0.6, 0.8 respectively. **b**-**d**. The performance (**b**), principal angle between rule subspaces (**c**) and the strength of conjunctive coding (**d**) before and after silencing SST neurons in the sensorimotor module, for networks trained with divisive dendritic nonlinearity (c.f. Figure 7e-g). **e**. The performance of many networks dropped to 0 when PV neurons in the sensorimotor module were silenced, indicating divergence in the network activity. **f**. The performance of many networks was not significantly changed when VIP neurons in the sensorimotor module were silenced. For **c**, One-sided Student’s t-test, *p* = 0.86, *n* = 24 networks. Each line represents a trained network. For **d**, One-sided Student’s t-test, *p* = 1.3 ∗ 10^−77^, *n* = 3430 units. Each line represents one unit. Results are aggregated across networks).

**Supplementary Figure 12:**
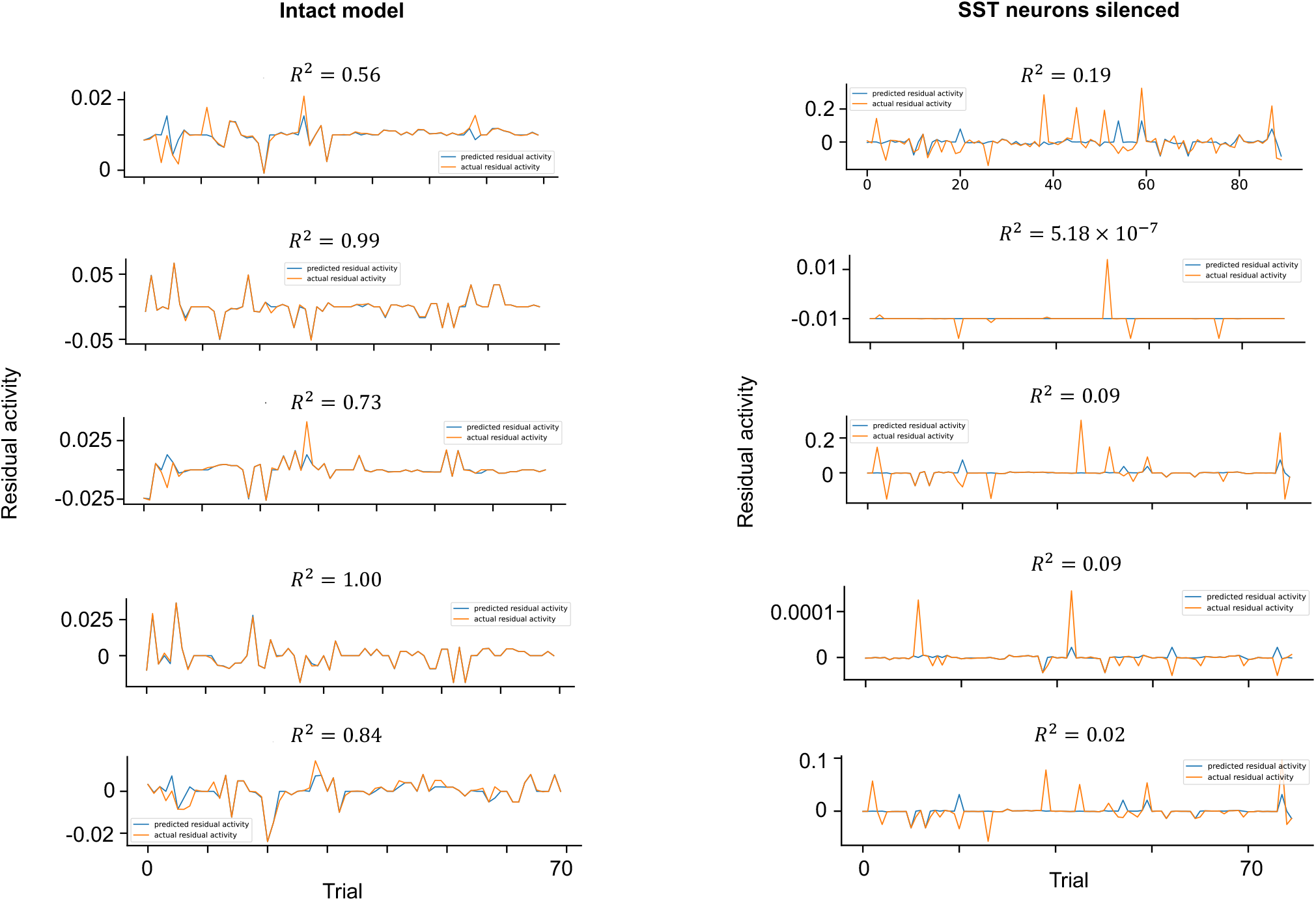
Example neurons in the sensorimotor module showing decreased conjunctive coding of rule and stimulus when SST neurons in the sensorimotor module are silenced. The strength of conjunctive coding is assessed by the *R*^2^ value of a linear regression model where the independent variables are conjunctions of rule and stimulus and the dependent variable is the trial-to-trial residual neural activity unexplained by a model with only rule and stimulus as independent variables (see Methods for details).

## References

[1] David A Grant and Esta Berg. A behavioral analysis of degree of reinforcement and ease of shifting to new responses in a weigl-type card-sorting problem. Journal of experimental psychology, 38(4):404, 1948.

[2] Valerio Mante, David Sussillo, Krishna V Shenoy, and William T Newsome. Context-dependent computation by recurrent dynamics in prefrontal cortex. nature, 503(7474):78–84, 2013.

[3] Rajeev V Rikhye, Aditya Gilra, and Michael M Halassa. Thalamic regulation of switching between cortical representations enables cognitive flexibility. Nature neuro-science, 21(12):1753–1763, 2018.

[4] Brenda Milner. Effects of different brain lesions on card sorting: The role of the frontal lobes. Archives of neurology, 9(1):90–100, 1963.

[5] R Dias, TW Robbins, and AC Roberts. Primate analogue of the wisconsin card sorting test: effects of excitotoxic lesions of the prefrontal cortex in the marmoset. Behavioral neuroscience, 110:872, 1996.

[6] RE Passingham. Non-reversal shifts after selective prefrontal ablations in monkeys (macaca mulatta). Neuropsychologia, 10(1):41–46, 1972.

[7] Katsuyuki Sakai. Task set and prefrontal cortex. Annu. Rev. Neurosci., 31:219–245, 2008.

[8] M. J. Buckley, F. A. Mansouri, H. Hoda, M. Mahboubi, P. G. Browning, S. C. Kwok, A. Phillips, and K. Tanaka. Dissociable components of rule-guided behavior depend on distinct medial and prefrontal regions. Science, 325:52–58, 2009.

[9] E. K. Miller and J. D. Cohen. An integrative theory of prefrontal cortex function. Annual Review of Neuroscience, 24:167–202, 2001.

[10] Farshad A Mansouri, Kenji Matsumoto, and Keiji Tanaka. Prefrontal cell activities related to monkeys’ success and failure in adapting to rule changes in a wisconsin card sorting test analog. Journal of Neuroscience, 26(10):2745–2756, 2006.

[11] Tsukasa Kamigaki, Tetsuya Fukushima, and Yasushi Miyashita. Cognitive set recon-figuration signaled by macaque posterior parietal neurons. Neuron, 61(6):941–951, 2009.

[12] Morteza Sarafyazd and Mehrdad Jazayeri. Hierarchical reasoning by neural circuits in the frontal cortex. Science, 364(6441):eaav8911, 2019.

[13] Takuya Ito, Guangyu Robert Yang, Patryk Laurent, Douglas H Schultz, and Michael W Cole. Constructing neural network models from brain data reveals representational transformations linked to adaptive behavior. Nature communications, 13(1):673, 2022.

[14] Kristen Delevich, Jason Tucciarone, Z Josh Huang, and Bo Li. The mediodorsal thalamus drives feedforward inhibition in the anterior cingulate cortex via parvalbumin interneurons. Journal of Neuroscience, 35(14):5743–5753, 2015.

[15] Hyun-Jae Pi, Balázs Hangya, Duda Kvitsiani, Joshua I Sanders, Z Josh Huang, and Adam Kepecs. Cortical interneurons that specialize in disinhibitory control. Nature, 503(7477):521–524, 2013.

[16] Siyu Zhang, Min Xu, Tsukasa Kamigaki, Johnny Phong Hoang Do, Wei-Cheng Chang, Sean Jenvay, Kazunari Miyamichi, Liqun Luo, and Yang Dan. Long-range and local circuits for top-down modulation of visual cortex processing. Science, 345(6197):660–665, 2014.

[17] William Muñoz, Robin Tremblay, Daniel Levenstein, and Bernardo Rudy. Layer-specific modulation of neocortical dendritic inhibition during active wakefulness. Science, 355(6328):954–959, 2017.

[18] Andreas J Keller, Mario Dipoppa, Morgane M Roth, Matthew S Caudill, Alessandro Ingrosso, Kenneth D Miller, and Massimo Scanziani. A disinhibitory circuit for contextual modulation in primary visual cortex. Neuron, 108(6):1181–1193, 2020.

[19] X. J. Wang, J. Tegnér, C. Constantinidis, and P. S. Goldman-Rakic. Division of labor among distinct subtypes of inhibitory neurons in a cortical microcircuit of working memory. Proceedings of the National Academy of Science, USA, 101(5):1368–73, 2004.

[20] A. Kepecs and G. Fishell. Interneuron cell types are fit to function. Nature, 505:318–326, 2014.

[21] R. Tremblay, S. Lee, and B. Rudy. GABAergic interneurons in the neocortex: From cellular properties to circuits. Neuron, 91:260–292, 2016.

[22] Guangyu Robert Yang, John D Murray, and Xiao-Jing Wang. A dendritic disinhibitory circuit mechanism for pathway-specific gating. Nature communications, 7(1):1–14, 2016.

[23] Kiyoshi Nakahara, Toshihiro Hayashi, Seiki Konishi, and Yasushi Miyashita. Functional mri of macaque monkeys performing a cognitive set-shifting task. Science, 295(5559):1532–1536, 2002.

[24] G. R. Yang and X.-J. Wang. Artificial neural networks for neuroscientists: a primer. Neuron, 107:1048–1070, 2020.

[25] Monika Jadi, Alon Polsky, Jackie Schiller, and Bartlett W Mel. Location-dependent effects of inhibition on local spiking in pyramidal neuron dendrites. PLoS computational biology, 8(6):e1002550, 2012.

[26] Guangyu Robert Yang, Madhura R Joglekar, H Francis Song, William T Newsome, and Xiao-Jing Wang. Task representations in neural networks trained to perform many cognitive tasks. Nature neuroscience, 22(2):297–306, 2019.

[27] H Francis Song, Guangyu R Yang, and Xiao-Jing Wang. Training excitatory-inhibitory recurrent neural networks for cognitive tasks: a simple and flexible frame-work. PLoS computational biology, 12(2):e1004792, 2016.

[28] Tsukasa Kamigaki, Tetsuya Fukushima, and Yasushi Miyashita. Neuronal signal dynamics during preparation and execution for behavioral shifting in macaque posterior parietal cortex. Journal of cognitive neuroscience, 23(9):2503–2520, 2011.

[29] Joaquin M Fuster. Unit activity in prefrontal cortex during delayed-response performance: neuronal correlates of transient memory. Journal of neurophysiology, 36(1):61–78, 1973.

[30] Shintaro Funahashi, Charles J Bruce, and Patricia S Goldman-Rakic. Mnemonic coding of visual space in the monkey’s dorsolateral prefrontal cortex. Journal of neurophysiology, 61(2):331–349, 1989.

[31] P. Goldman-Rakic. Cellular basis of working memory. Neuron, 14:477–85, 1995.

[32] Ranulfo Romo, Carlos D Brody, Adrián Hernández, and Luis Lemus. Neuronal correlates of parametric working memory in the prefrontal cortex. Nature, 399(6735):470–473, 1999.

[33] Thomas B Christophel, P Christiaan Klink, Bernhard Spitzer, Pieter R Roelfsema, and John-Dylan Haynes. The distributed nature of working memory. Trends in cognitive sciences, 21(2):111–124, 2017.

[34] Matthew L Leavitt, Diego Mendoza-Halliday, and Julio C Martinez-Trujillo. Sustained activity encoding working memories: not fully distributed. Trends in Neurosciences, 40(6):328–346, 2017.

[35] Zengcai V Guo, Hidehiko K Inagaki, Kayvon Daie, Shaul Druckmann, Charles R Gerfen, and Karel Svoboda. Maintenance of persistent activity in a frontal thalamo-cortical loop. Nature, 545(7653):181–186, 2017.

[36] Kartik K Sreenivasan and Mark D’Esposito. The what, where and how of delay activity. Nature reviews neuroscience, 20(8):466–481, 2019.

[37] Kong-Fatt Wong and Xiao-Jing Wang. A recurrent network mechanism of time integration in perceptual decisions. Journal of Neuroscience, 26(4):1314–1328, 2006.

[38] Farzaneh Najafi, Gamaleldin F Elsayed, Robin Cao, Eftychios Pnevmatikakis, Peter E Latham, John P Cunningham, and Anne K Churchland. Excitatory and inhibitory subnetworks are equally selective during decision-making and emerge simultaneously during learning. Neuron, 105(1):165–179, 2020.

[39] James P Roach, Anne K Churchland, and Tatiana A Engel. Choice selective inhibition drives stability and competition in decision circuits. Nature Communications, 14(1):147, 2023.

[40] Hongbo Jia, Nathalie L Rochefort, Xiaowei Chen, and Arthur Konnerth. Dendritic organization of sensory input to cortical neurons in vivo. Nature, 464(7293):1307–1312, 2010.

[41] Joseph Cichon and Wen-Biao Gan. Branch-specific dendritic ca2+ spikes cause persistent synaptic plasticity. Nature, 520(7546):180–185, 2015.

[42] Shannon K Rashid, Victor Pedrosa, Martial A Dufour, Jason J Moore, Spyridon Chavlis, Rodrigo G Delatorre, Panayiota Poirazi, Claudia Clopath, and Jayeeta Basu. The dendritic spatial code: branch-specific place tuning and its experience-dependent decoupling. BioRxiv, pages 2020–01, 2020.

[43] Jakob Voigts and Mark T Harnett. Somatic and dendritic encoding of spatial variables in retrosplenial cortex differs during 2d navigation. Neuron, 105(2):237–245, 2020.

[44] Mattia Rigotti, Omri Barak, Melissa R Warden, Xiao-Jing Wang, Nathaniel D Daw, Earl K Miller, and Stefano Fusi. The importance of mixed selectivity in complex cognitive tasks. Nature, 497(7451):585–90, 2013.

[45] Atsushi Kikumoto and Ulrich Mayr. Conjunctive representations that integrate stimuli, responses, and rules are critical for action selection. Proceedings of the National Academy of Sciences, 117(19):10603–10608, 2020.

[46] Atsushi Kikumoto, Ulrich Mayr, and David Badre. The role of conjunctive representations in prioritizing and selecting planned actions. Elife, 11:e80153, 2022.

[47] Jonathan D Wallis, Kathleen C Anderson, and Earl K Miller. Single neurons in prefrontal cortex encode abstract rules. Nature, 411(6840):953–956, 2001.

[48] Matthew M Botvinick, Jonathan D Cohen, and Cameron S Carter. Conflict monitoring and anterior cingulate cortex: an update. Trends in cognitive sciences, 8(12):539–546, 2004.

[49] René Quilodran Marie Rothe, and Emmanuel Procyk. Behavioral shifts and action valuation in the anterior cingulate cortex. Neuron, 57(2):314–325, 2008.

[50] Nils Kolling, Marco K Wittmann, Tim EJ Behrens, Erie D Boorman, Rogier B Mars, and Matthew FS Rushworth. Value, search, persistence and model updating in anterior cingulate cortex. Nature neuroscience, 19(10):1280–1285, 2016.

[51] Farshad Alizadeh Mansouri, David J Freedman, and Mark J Buckley. Emergence of abstract rules in the primate brain. Nature Reviews Neuroscience, 21(11):595–610, 2020.

[52] Timothy Spellman, Malka Svei, Jesse Kaminsky, Gabriela Manzano-Nieves, and Conor Liston. Prefrontal deep projection neurons enable cognitive flexibility via persistent feedback monitoring. Cell, 184(10):2750–2766, 2021.

[53] C Daniel Salzman and Stefano Fusi. Emotion, cognition, and mental state representation in amygdala and prefrontal cortex. Annual review of neuroscience, 33:173–202, 2010.

[54] Clay B Holroyd and Michael GH Coles. The neural basis of human error processing: reinforcement learning, dopamine, and the error-related negativity. Psychological review, 109(4):679, 2002.

[55] Masayuki Matsumoto and Okihide Hikosaka. Two types of dopamine neuron distinctly convey positive and negative motivational signals. Nature, 459(7248):837–841, 2009.

[56] Richard A Andersen and He Cui. Intention, action planning, and decision making in parietal-frontal circuits. Neuron, 63(5):568–583, 2009.

[57] Bernard W Balleine and John P O’doherty. Human and rodent homologies in action control: corticostriatal determinants of goal-directed and habitual action. Neuropsychopharmacology, 35(1):48–69, 2010.

[58] Bevil R Conway. Color vision, cones, and color-coding in the cortex. The neuroscientist, 15(3):274–290, 2009.

[59] Rosa Lafer-Sousa and Bevil R Conway. Parallel, multi-stage processing of colors, faces and shapes in macaque inferior temporal cortex. Nature neuroscience, 16(12):1870–1878, 2013.

[60] Le Chang, Pinglei Bao, and Doris Y Tsao. The representation of colored objects in macaque color patches. Nature communications, 8(1):2064, 2017.

[61] Mark M Churchland, John P Cunningham, Matthew T Kaufman, Justin D Foster, Paul Nuyujukian, Stephen I Ryu, and Krishna V Shenoy. Neural population dynamics during reaching. Nature, 487(7405):51–56, 2012.

[62] Nicholas A Steinmetz, Peter Zatka-Haas, Matteo Carandini, and Kenneth D Harris. Distributed coding of choice, action and engagement across the mouse brain. Nature, 576(7786):266–273, 2019.

[63] Shih-Yi Tseng, Selmaan N Chettih, Charlotte Arlt, Roberto Barroso-Luque, and Christopher D Harvey. Shared and specialized coding across posterior cortical areas for dynamic navigation decisions. Neuron, 110(15):2484–2502, 2022.

[64] Charles Findling, Felix Hubert, International Brain Laboratory, Luigi Acerbi, Brandon Benson, Julius Benson, Daniel Birman, Niccolò Bonacchi Matteo Carandini, Joana A Catarino, et al. Brain-wide representations of prior information in mouse decision-making. BioRxiv, pages 2023–07, 2023.

[65] International Brain Lab, Brandon Benson, Julius Benson, Daniel Birman, Niccolo Bonacchi, Matteo Carandini, Joana A Catarino, Gaelle A Chapuis, Anne K Churchland, Yang Dan, et al. A brain-wide map of neural activity during complex behaviour. bioRxiv, pages 2023–07, 2023.

[66] Sean Froudist-Walsh, Daniel P Bliss, Xingyu Ding, Lucija Rapan, Meiqi Niu, Kenneth Knoblauch, Karl Zilles, Henry Kennedy, Nicola Palomero-Gallagher, and Xiao-Jing Wang. A dopamine gradient controls access to distributed working memory in the large-scale monkey cortex. Neuron, 109(21):3500–3520, 2021.

[67] Jorge F Mejías and Xiao-Jing Wang. Mechanisms of distributed working memory in a large-scale network of macaque neocortex. Elife, 11:e72136, 2022.

[68] Mattia Rigotti, Daniel Ben Dayan Rubin, Xiao-Jing Wang, and Stefano Fusi. Internal representation of task rules by recurrent dynamics: the importance of the diversity of neural responses. Frontiers in computational neuroscience, 4:24, 2010.

[69] Stanislas Dehaene and Jean-Pierre Changeux. The wisconsin card sorting test: Theoretical analysis and modeling in a neuronal network. Cerebral cortex, 1(1):62–79, 1991.

[70] Daniel Turner-Evans, Stephanie Wegener, Herve Rouault, Romain Franconville, Tanya Wolff, Johannes D Seelig, Shaul Druckmann, and Vivek Jayaraman. Angular velocity integration in a fly heading circuit. Elife, 6:e23496, 2017.

[71] Benjamin Y Hayden, John M Pearson, and Michael L Platt. Neuronal basis of sequential foraging decisions in a patchy environment. Nature neuroscience, 14(7):933–939, 2011.

[72] João D Semedo, Amin Zandvakili, Christian K Machens, M Yu Byron, and Adam Kohn. Cortical areas interact through a communication subspace. Neuron, 102(1):249–259, 2019.

[73] Christopher Langdon, Mikhail Genkin, and Tatiana A Engel. A unifying perspective on neural manifolds and circuits for cognition. Nature Reviews Neuroscience, pages 1–15, 2023.

[74] Seung-Hee Lee, Alex C Kwan, Siyu Zhang, Victoria Phoumthipphavong, John G Flannery, Sotiris C Masmanidis, Hiroki Taniguchi, Z Josh Huang, Feng Zhang, Edward S Boyden, et al. Activation of specific interneurons improves v1 feature selectivity and visual perception. Nature, 488(7411):379–383, 2012.

[75] Seung-Hee Lee, Alex C Kwan, and Yang Dan. Interneuron subtypes and orientation tuning. Nature, 508(7494):E1–E2, 2014.

[76] Nathan R Wilson, Caroline A Runyan, Forea L Wang, and Mriganka Sur. Division and subtraction by distinct cortical inhibitory networks in vivo. Nature, 488(7411):343–348, 2012.

[77] Yu Fu, Jason M Tucciarone, J Sebastian Espinosa, Nengyin Sheng, Daniel P Darcy, Roger A Nicoll, Z Josh Huang, and Michael P Stryker. A cortical circuit for gain control by behavioral state. Cell, 156(6):1139–1152, 2014.

[78] Carsten K Pfeffer, Mingshan Xue, Miao He, Z Josh Huang, and Massimo Scanziani. Inhibition of inhibition in visual cortex: the logic of connections between molecularly distinct interneurons. Nature neuroscience, 16(8):1068–1076, 2013.

[79] Sahil Loomba, Jakob Straehle, Vijayan Gangadharan, Natalie Heike, Abdelrahman Khalifa, Alessandro Motta, Niansheng Ju, Meike Sievers, Jens Gempt, Hanno S Meyer, et al. Connectomic comparison of mouse and human cortex. Science, 377(6602):eabo0924, 2022.

[80] Braden A Purcell and Roozbeh Kiani. Hierarchical decision processes that operate over distinct timescales underlie choice and changes in strategy. Proceedings of the national academy of sciences, 113(31):E4531–E4540, 2016.

[81] Kevin Johnston, Helen M Levin, Michael J Koval, and Stefan Everling. Top-down control-signal dynamics in anterior cingulate and prefrontal cortex neurons following task switching. Neuron, 53(3):453–462, 2007.

[82] Ken-Ichiro Tsutsui, Takayuki Hosokawa, Munekazu Yamada, and Toshio Iijima. Representation of functional category in the monkey prefrontal cortex and its ruledependent use for behavioral selection. Journal of Neuroscience, 36(10):3038–3048, 2016.

[83] Silvia Bernardi, Marcus K Benna, Mattia Rigotti, Jérôme Munuera, Stefano Fusi, and C Daniel Salzman. The geometry of abstraction in the hippocampus and prefrontal cortex. Cell, 183(4):954–967, 2020.

[84] Vishwa Goudar, Jeong-Woo Kim, Yue Liu, Adam JO Dede, Michael J Jutras, Ivan Skelin, Michael Ruvalcaba, William Chang, Adrienne L Fairhall, Jack J Lin, et al. Comparing rapid rule-learning strategies in humans and monkeys. bioRxiv, pages 2023–01, 2023.

[85] Steven W Kennerley, Mark E Walton, Timothy EJ Behrens, Mark J Buckley, and Matthew FS Rushworth. Optimal decision making and the anterior cingulate cortex. Nature neuroscience, 9(7):940–947, 2006.

[86] Cheng Xue, Lily E Kramer, and Marlene R Cohen. Dynamic task-belief is an integral part of decision-making. Neuron, 110(15):2503–2511, 2022.

[87] Ido Ben-Artzi, Yoav Kessler, Bruno Nicenboim, and Nitzan Shahar. Computational mechanisms underlying latent value updating of unchosen actions. Science Advances, 9(42):eadi2704, 2023.

[88] Amitai Shenhav, Matthew M Botvinick, and Jonathan D Cohen. The expected value of control: an integrative theory of anterior cingulate cortex function. Neuron, 79(2):217–240, 2013.

[89] Stefano Fusi, Wael F Asaad, Earl K Miller, and Xiao-Jing Wang. A neural circuit model of flexible sensorimotor mapping: learning and forgetting on multiple timescales. Neuron, 54(2):319–333, 2007.

[90] Paul G Anastasiades, David P Collins, and Adam G Carter. Mediodorsal and ventromedial thalamus engage distinct l1 circuits in the prefrontal cortex. Neuron, 109(2):314–330, 2021.

[91] Arghya Mukherjee, Norman H Lam, Ralf D Wimmer, and Michael M Halassa. Thalamic circuits for independent control of prefrontal signal and noise. Nature, 600(7887):100–104, 2021.

[92] Pablo Garcia-Junco-Clemente, Taruna Ikrar, Elaine Tring, Xiangmin Xu, Dario L Ringach, and Joshua T Trachtenberg. An inhibitory pull–push circuit in frontal cortex. Nature neuroscience, 20(3):389–392, 2017.

[93] Zhixiao Su and Jeremiah Y Cohen. Two types of locus coeruleus norepinephrine neurons drive reinforcement learning. bioRxiv, pages 2022–12, 2022.

[94] Jounhong Ryan Cho, Jennifer B Treweek, J Elliott Robinson, Cheng Xiao, Lindsay R Bremner, Alon Greenbaum, and Viviana Gradinaru. Dorsal raphe dopamine neurons modulate arousal and promote wakefulness by salient stimuli. Neuron, 94(6):1205–1219, 2017.

[95] Xiaolong Jiang, Shan Shen, Cathryn R Cadwell, Philipp Berens, Fabian Sinz, Alexander S Ecker, Saumil Patel, and Andreas S Tolias. Principles of connectivity among morphologically defined cell types in adult neocortex. Science, 350(6264):aac9462, 2015.

[96] Paul J Werbos. Backpropagation through time: what it does and how to do it. Proceedings of the IEEE, 78(10):1550–1560, 1990.

[97] Diederik P Kingma and Jimmy Ba. Adam: A method for stochastic optimization. arXiv preprint 1412.6980, 2014.

[98] Camille Jordan. Essai sur la géométrie à n dimensions. Bulletin de la Société mathématique de France, 3:103–174, 1875.

[99] ke Björck and Gene H Golub. Numerical methods for computing angles between linear subspaces. Mathematics of computation, 27(123):579–594, 1973.

[100] Peiran Gao, Eric Trautmann, Byron Yu, Gopal Santhanam, Stephen Ryu, Krishna Shenoy, and Surya Ganguli. A theory of multineuronal dimensionality, dynamics and measurement. BioRxiv, page 214262, 2017.

[101] Yue Liu. Flexible gating between subspaces in a neural network model of internally guided task switching.

